# Stimulation with ECoG electrodes modulates cortical activity and sensory processing in the awake mouse brain

**DOI:** 10.1101/2025.10.28.685211

**Authors:** Jiang Lan Fan, Keundong Lee, Youngbin Tchoe, Mehran Ganji, Ritwik Vatsyayan, Ha Yun Yoon, Jacob Garrett, Shadi A. Dayeh, Eric Halgren, Na Ji

**Affiliations:** Joint Bioengineering Graduate Program, University of California, Berkeley, and University of California, San Francisco, CA, USA; Department of Electrical and Computer Engineering, University of California, San Diego, La Jolla, CA 92093, USA; Department of Mechanical Engineering, University of California, Berkeley, CA, 97420, USA; Departments of Radiology and Neurosciences, University of California, San Diego, CA 92093, USA; Department of Physics, University of California, Berkeley, CA, 94720, USA; Department of Neuroscience, University of California, Berkeley, CA, 94720, USA; Helen Wills Neuroscience Institute, University of California, Berkeley, CA, 94720, USA; Molecular Biophysics and Integrated Bioimaging Division, Lawrence Berkeley National Laboratory, Berkeley, CA, 94720, USA

## Abstract

Electrical stimulation has been widely used to probe neural network properties and treat dysfunction. Electrocorticography (ECoG) electrodes, long used for activity monitoring, can also stimulate the brain in a minimally invasive and chronic manner. However, how cortical surface electrical stimulation impacts cortical network activity remains poorly understood. Using in vivo calcium imaging in the awake mouse brain with chronically implanted ECoG electrodes, we measured how electrical stimulation modulates the activity of visual cortical neurons, including during concurrent visual stimulation. We found that cortical surface electrical stimulation initially activates L2/3 neurons followed by prolonged inhibition lasting seconds after stimulation. Electrical stimulation suppresses the activity of neurons at their preferred grating orientation but enhances their responses to non-preferred visual stimuli, thereby reducing sensory feature selectivity. By measuring how electrical stimulation modulates the activity of inhibitory neuron subtypes including PV, SST, and NDNF interneurons, we propose a circuit model in which L1 NDNF interneurons are strongly activated by cortical electrical stimulation and, in turn, inhibit L2/3 excitatory neurons and PV interneurons through volume transmission of GABA.

## Introduction

Electrocorticography (ECoG), which employs a flexible sheet of electrode contacts placed on the surface of the brain, has been widely used in clinical settings to record electrical activity for epilepsy monitoring and for mapping eloquent cortex, thereby allowing the resection of tumors, epileptic foci, or other pathology without causing unacceptable functional deficits^1–3^. Cortical mapping with ECoG is performed using electrical stimulation to evoke sensation or movement, or to disrupt language task performance. In addition to providing essential clinical information, it has fundamentally shaped our concepts of human cortical functional organization^4–6^. ECoG stimulation is currently being explored for brain-computer interface (BCI) applications, including sensory and motor prostheses, as well as modulation of depression and pain^7–11^. Despite its clinical and experimental importance, the circuit mechanisms through which ECoG stimulation modulates cortical neuronal activity are poorly understood, hindering efforts to optimize ECoG array design and stimulation protocols for both basic research and clinical applications.

This gap in knowledge reflects a broader lack of understanding of how electrical stimulation affects neuronal activity, especially at the circuit level and with cell-type specificity, despite its longstanding use to probe brain function^12–14^ and treat neurological disorders^15–19^. Most prior investigations have focused on intracortical electrical stimulation. A seminal study from 1968 concluded that stimulation above certain current thresholds directly excites the somata of pyramidal (PYR) neurons within a spherical region around the electrode tip, with the size of the sphere increasing with stimulation current strength^20^. In contrast, a 2009 study using calcium indicators and two-photon fluorescence microscopy (2PFM)^21^ observed spatially sparse and distributed activation of neurons. Challenging the earlier model, it concluded that intracortical stimulation sparsely activates nearby neuronal cell bodies but directly activates nearby axons. Through antidromic activation of distally projecting axons, electrical stimulation can excite neurons located millimeters away. This is consistent with previous work identifying the nodes of Ranvier and axon hillocks as the sites with the lowest thresholds for externally imposed current gradients to trigger action potentials^22–26^. Additional mechanisms come into play when stimulation is applied in trains^25,27,28^ or occurs in the context of pre-stimulus activity^29^, resulting in a complex recruitment of both PYR and inhibitory neurons (INs). These considerations have proven important for understanding the mechanism of the most common BCI therapy of deep brain stimulation for Parkinson’s disease^30^.

In this study, we investigated how cortical surface stimulation using ECoG electrodes modulates cortical neuron activity in the awake mouse cortex using 2PFM and cell-type specific expression of calcium indicator GCaMP6s. We further explored how electrical stimulation affects cortical sensory processing by combining electrical and visual stimulation. We found that cortical surface stimulation alone first directly activates L2/3 PYR neurons and L1 NDNF INs, then suppresses their ongoing activity to below baseline level. Electrical stimulation interacts with visual stimulation additively when neurons are weakly excited by visual stimulation, but suppresses the activity of neurons exhibiting strong visually evoked activity in a divisive manner, leading to an overall reduction in sensory feature selectivity. Whereas PV and SST IN activity are moderately impacted by electrical stimulation, L1 NDNF neurons appear to play a major role in shaping cortical circuit activity by mediating a sustained suppression of ongoing and sensory-evoked activity.

We propose a circuit model in which cortical surface stimulation directly activates L2/3 PYR and NDNF INs through antidromic activation of their axon projections in L1, leading to a rapid increase in their firing rates. The strong recruitment of NDNF inhibitory neurons in turn causes a strong and sustained inhibition of L2/3 PYR as well as NDNF and PV IN populations, likely through GABA volume transmission. Our results indicate that the composition, geometrical arrangement, electrical properties, and connectivity of excitatory and inhibitory subtypes near the ECoG electrodes dictate how electrical stimulation modulates neuronal responses and impacts information processing in the cortex.

## Results

### A novel cortical surface ECoG implant enables concurrent electrical stimulation and 2PFM imaging in the awake mouse brain

We developed a chronic implant that allowed simultaneous microstimulation and 2PFM imaging of the awake mouse cortex (**Supplementary Fig. 1**, Methods). A surface electrode array resembling a miniaturized human ECoG array was fabricated by developing platinum nanorod (PtNR) films on electrode contact surfaces. The resulting round PtNR electrodes were 90 or 200 µm in diameter. Glued to a glass cranial window, the electrode array was implanted during a craniotomy surgery above the mouse’s left primary visual cortex (V1) with its electrodes in contact with the intact dura and had <60 kΩ impedances at 1 kHz measured post implantation. The cranial window provided mechanical support to the electrode array as well as chronic optical access to the brain tissue below. During the same surgery, AAV2/1 viral particles were injected into the cortex to drive expression of the genetically encoded calcium indicator GCaMP6s^31^, following previously described procedure^32,33^. A stainless-steel head-bar was attached to the skull and a 3D-printed resin housing was attached to the head-bar. The housing shielded stray light during imaging and protected the electrode-bonded PCB.

We carried out implant surgeries and expressed GCaMP6s in 14 mice, including wildtype mice and three transgenic lines that allowed Cre-dependent GCaMP6s expression in PV, SOM, and NDNF subtypes of inhibitory neurons^34–37^, respectively. After 2 weeks of viral transduction and habituation for head fixation, we imaged GCaMP6s+ V1 neurons in the head-fixed awake mouse at 15 frames/s using a previously described 2PFM system^32^ operated either in the standard Gaussian focus scanning mode or, to increase imaging throughput, the Bessel focus scanning mode^38,39^. A computer monitor was used to present drifting grating visual stimuli to the right eye of the animal (**Fig. 1a**). Imaging fields of view (FOVs), typically 781 µm by 781 µm, were chosen within V1 by retinotopic mapping of 2.2 mm × 2.2 mm cortical areas (Methods). The same FOVs were then imaged during symmetric biphasic pulse stimulation by a nearby electrode and/or visual stimulation (**Fig 1b,c**).

**Figure 1.**
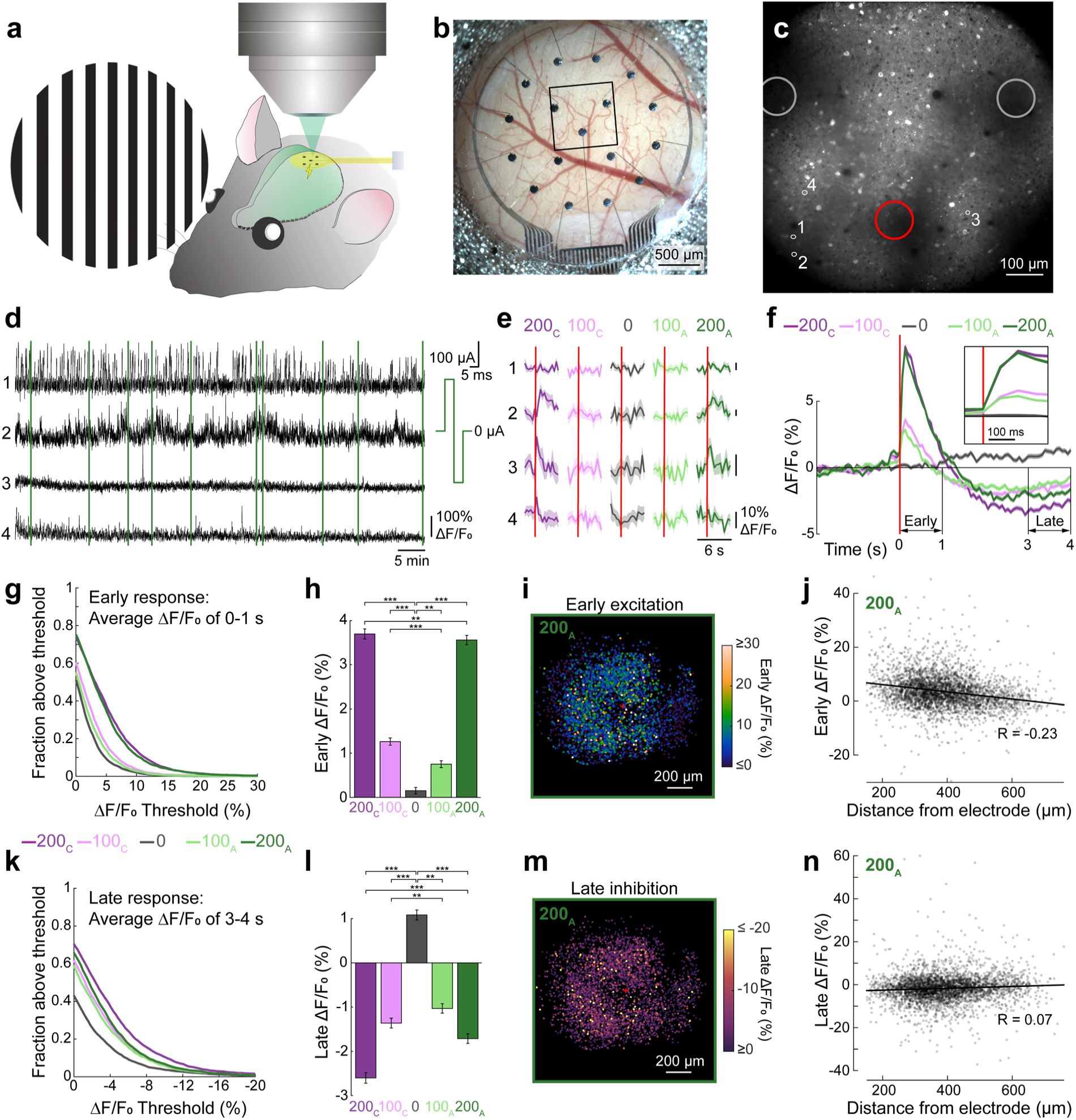
Cortical-surface electrical stimulation first activates then suppresses L2/3 neuron activity in the mouse V1. (**a**) Setup for concurrent 2PFM, electrical stimulation (E-Stim), and visual stimulation of the awake mouse V1. (**b**) Image of a cranial window and ECoG electrode 2 weeks post implant. (**c**) 2PFM image of GCaMP6s+ neurons 200 µm below dura within the black box in **b** (displayed at γ=0.5). Large circles: profile of electrodes. Red circle: stimulation electrode. (**d**) Calcium activity ΔF/F_0_ traces of example ROI 1-4 outlined in **c** over 75 min. Green lines: pseudorandomly-distributed 10 trials of 200 µA anode-first stimulation (inset, 200_A_) without visual stimulation. (**e**) Trial-averaged responses of the ROIs in **d** to electrical stimuli. Trace and shade: mean and SEM. Red lines: onset of E-Stim. (**f**) Average ΔF/F_0_ traces of 3570 L2/3 neurons in response to E-Stim (red line). Trace and shade: mean and SEM. Inset: zoomed-in to 0-200 ms post E-Stim. (**g**) Fractions of neurons with their early response (average ΔF/F_0_ during 0-1 s post E-Stim) above a specific threshold value vs. threshold values, for all stimulation currents. (**h**) Early responses of 3570 neurons. Error bars: SEM. Two-sample KS-test; p values: *: < 0.05, **: < 0.01, ***: < 0.001. (**i**) Early responses to 200_A_ E-Stim, plotted for each neuron based on their lateral displacement from stimulation electrode (red “+”: electrode center). (**j**) Early response to 200_A_ E-Stim of each neuron vs. its 3D distance from electrode center. R: Pearson’s correlation coefficient. (**k,l,m,n**) Same as (**g,h,i,j**) but for late responses (time-averaged ΔF/F_0_ during 3-4 s post E-Stim) to 200_A_ E-Stim.

We evaluated the calcium responses of the same neurons to electrical stimulus alone (E-Stim), visual stimulus alone (V-Stim), and concurrent electrical and visual stimulation (EV-Stim), respectively. With 5 electrical stimulation settings (a single 10-ms-long symmetric biphasic pulse with currents at −200 µA, −100 µA, 0, 100 µA, or 200 µA) and 9 visual stimulation types (no visual stimulation, 8 gratings of spatial frequency of 0.07 cycles/degree and temporal frequency of 2 cycles/second, drifting along 0°, 45°, 90°, 135°, 180°, 225°, 270°, 315° directions), there were a total of 45 unique stimulation conditions. With each condition repeated for 10 trials, an imaging session included 450 trials ordered pseudo-randomly, with each trial composed of a 2-s baseline measurement, onset of stimulation, and 4-s measurement of stimulation response. There was a 4-s gap in image acquisition between trials to allow the calcium response evoked in the previous trial to decay back to baseline. After image registration, the fluorescence trace F of each hand-segmented soma was extracted and the neuropil background subtracted (Methods). Calcium activity trace ΔF/F_0_ (F_0_: baseline fluorescence, calculated from the 2-s baseline measurement; ΔF = F - F_0_) was calculated for each soma (e.g., example neurons 1-4, **Fig. 1d**) and a neuron’s response to each unique stimulus was calculated as the averaged ΔF/F_0_ of 10 trials (e.g., responses to E-Stim by neurons 1-4, **Fig. 1e**). Statistical tests were used to determine the response properties of each neuron (Methods).

### Cortical surface electrical stimulation first activates then suppresses L2/3 population activity in the mouse V1

We first evaluated the responses of L2/3 neurons (150-270 µm below dura) evoked by cortical surface electrical stimulation in the absence of visual stimulation. ΔF/F_0_ traces of four representative neurons in a wildtype mouse V1 are shown with the ten anode-leading 200 µA stimulation events marked with vertical green lines (**Fig. 1d**). As indicated by the trial-averaged responses of these neurons to 10-ms-long electrical stimulation of cathode-leading 200 µA (“200_C_”), cathode-leading 100 µA (“100_C_”), 0, anode-leading 100 µA (“100_A_”), or anode-leading 200 µA (“200_A_”) symmetric biphasic current injections (**Fig. 1e**), some neurons exhibited electrically evoked responses (e.g., Neurons 2-4) while others did not (e.g., Neuron 1).

To investigate the population response of V1 neurons, we calculated the mean E-Stim-evoked calcium response of 3,570 L2/3 neurons in wildtype mice (3 animals, 14 FOVs; **Fig. 1f**). Immediately after electrical stimulation (onset: red line, **Fig. 1f**), we observed a rapid rise in calcium within the first frame (i.e., 67 ms post stimulation onset), peaking at the second frame (i.e., 133 ms post stimulation onset) (**Fig. 1f**, inset), with stronger stimulation currents evoking calcium transients of larger ΔF/F_0_.

Consistent with the mean E-Stim-evoked population calcium responses, more neurons had average ΔF/F_0_ during 0-1 s post E-Stim above threshold values when the stimulus current increased from 100 µA to 200 µA (**Fig. 1g**), indicating that larger currents excite higher fractions of neurons. Statistically, the averaged ΔF/F_0_ between 0 and 1 s after E-Stim onset, hereafter defined as the “early response”, of 3,570 neurons was significantly greater with electrical stimulation than without (**Fig. 1h**; two-sample Kolmogorov–Smirnov test; p values in **Supplementary Table 1**). A 200 µA stimulation current induced significantly larger fluorescence responses compared to 100 µA and cathode-leading pulses induced slightly larger fluorescence responses than anode-leading pulses (**Fig. 1h**; two-sample Kolmogorov– Smirnov test; p values: **Supplementary Table 1**, 200_C_ vs. 100_C_: 2.3×10^−77^, 200_A_ vs. 100_A_: 2.6×10^−101^).

Plotting the early response strengths of individual neurons relative to their 3D distances from the center of the stimulating electrode, we observed a weak negative correlation, i.e., stronger activation for neurons closer to the electrode, across all stimulation conditions (200_A_ data shown in **Fig. 1i,j**; p = 2.2×10^−45^, two-tailed test for Pearson’s correlation coefficient; other conditions, **Supplementary Fig. 2a**). The dependence of L2/3 neurons’ early responses on stimulation strength (with little or no change in waveform) and distance from electrode (especially compared to late responses, see below) is consistent with their being directly evoked by the electrical stimulation, as opposed to consequent network activity.

Interestingly, after the early activation response, ΔF/F_0_ reversed in sign and became negative (**Fig. 1f**), indicative of a strong late-onset inhibition of the population activity by the electrical stimulation, which reduced ongoing spontaneous activity. The inhibitory effect manifested itself strongly in the reduction of averaged ΔF/F_0_ between 3 and 4 s post E-Stim at both 100 µA and 200 µA stimulus current (**Fig. 1k**). The averaged ΔF/F_0_ between 3 and 4 s after E-Stim onset, hereafter defined as the “late response”, indicated that larger currents led to stronger inhibition (**Fig. 1l**; two-sample Kolmogorov–Smirnov test; p values: **Supplementary Table 2**, 200_A_ vs. 100_A_: 8.5×10^−13^, 200_C_ vs. 100_C_: 6.1×10^−19^). This late inhibition was less dependent on distance than the early excitation (200_A_, **Fig. 1m,n**; other conditions, **Supplementary Fig. 2b**). Its timing and lack of distance dependence suggest that this late onset inhibition was indirect in nature and involved the local cortical network.

### Electrical stimulation interacts with visual stimulation additively during weak visual responses and divisively during strong visual responses

Having found that cortical surface electrical stimulation has an initial activating effect followed by a suppressing effect on L2/3 population activity, we asked how electrical stimulation interacted with sensory processing by the same neurons. Neurons in L2/3 of the mouse V1 have visually evoked activity that can be selective for the orientation of drifting grating stimuli. Therefore, we measured and compared the calcium responses of L2/3 neurons in wildtype mice evoked by electrical stimulation alone (“E-Stim”), drifting grating visual stimulation alone (“V-Stim”), and simultaneous electrical and visual stimulation (“EV-Stim”).

For an example orientation-selective neuron, E-Stim evoked minimal calcium responses (column “No vis”, **Fig. 2a**), while V-Stim at its preferred grating evoked a calcium response of large ΔF/F_0_ magnitude (middle row, red arrow, **Fig. 2a**). Combining electrical and visual stimulation, instead of enhancing ΔF/F_0_ magnitude, substantially reduced the magnitudes of the visually evoked responses to the preferred drifting gratings (top two and bottom two rows, **Fig. 2a**). Fitting the averaged ΔF/F_0_ during the 4 seconds of visual stimulation across different drifting directions with a double Gaussian function for all electrical stimulation settings (**Fig. 2b**), we identified the preferred grating orientation of this neuron, which remained unchanged. Under EV-Stim, electrical stimulation reduced this neuron’s response to its preferred visual stimulus (red arrow, **Fig. 2b**), with larger currents leading to larger suppression.

**Figure 2.**
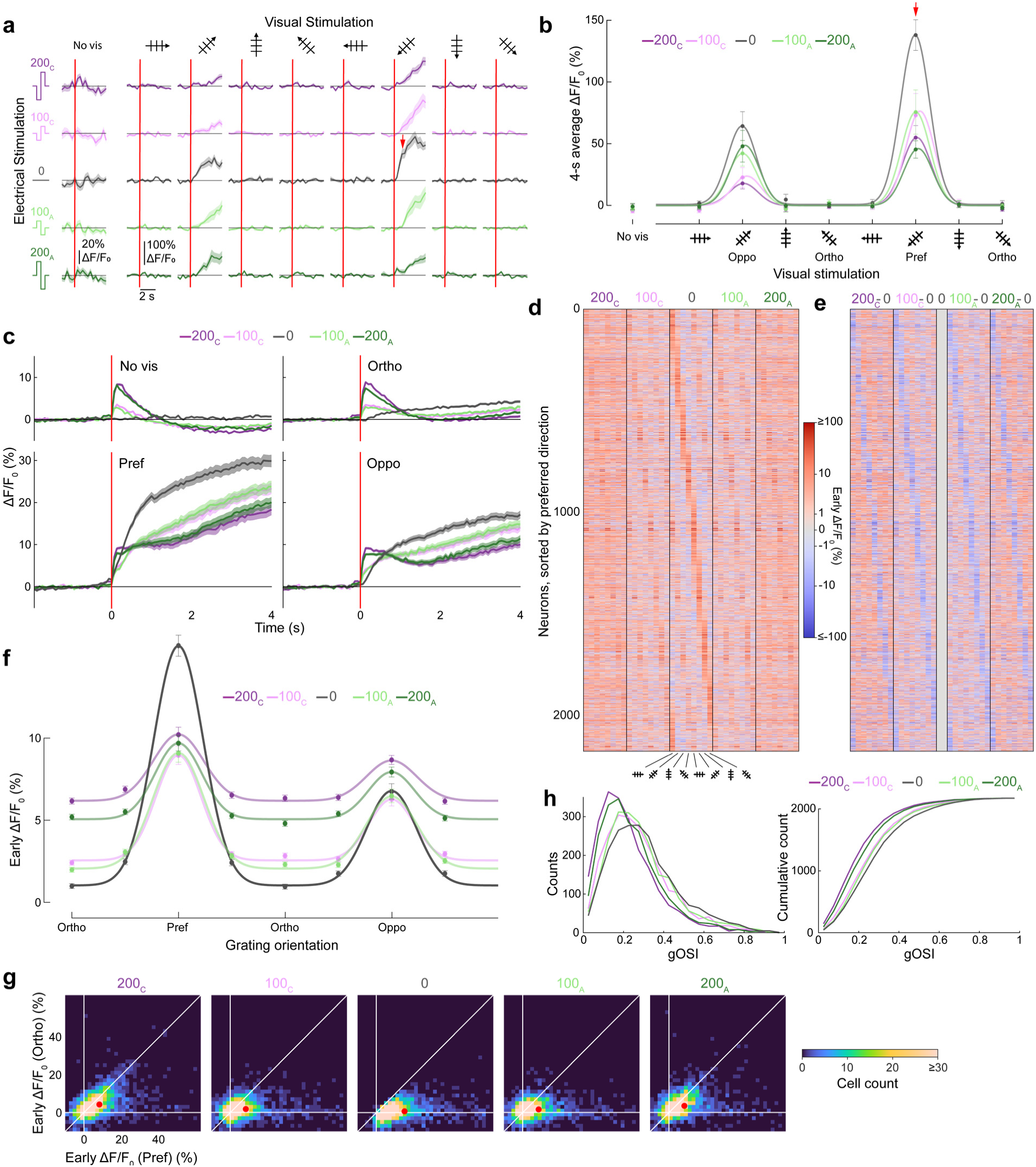
Electrical stimulation modulates visually evoked activity and reduces orientation selectivity in L2/3 neurons. (**a**) Trial-averaged ΔF/F_0_ traces of an example neuron responding to E-Stim, V-Stim, and EV-Stim. Trace and shade: mean and SEM. Red lines: stimulus onset. (**b**) Scattered data points: time-averaged ΔF/F_0_ over 4 s post stimulus onset of E-Stim, V-Stim, and EV-Stim; tuning curves: double-Gaussian fits to the responses towards 8 drifting gratings for the neuron in **a**. “No vis”: response to E-Stim; “oppo”, “ortho”, “pref”: response to V-Stim and EV-Stim for gratings moving in directions 180° to, ±90° to, and along the drifting direction of the preferred grating, respectively. (c) Population- and trial-averaged ΔF/F_0_ traces of 2173 L2/3 neurons with visually evoked activity under 5 electrical stimulation conditions, grouped for 4 visual stimulation conditions: no stimulus, ortho, pref, and oppo. Trace and shade: mean and SEM. Red lines: stimulus onset. (d) Heatmap showing the early response ΔF/F_0_ of 2173 L2/3 neurons. Rows: individual cells sorted by their preferred direction under V-Stim; Columns: electrical-visual stimulation conditions. (**e**) Heatmap showing the difference in early response ΔF/F_0_, by subtracting V-Stim responses from EV-Stim responses in **d**. Blue: reduced activity; red: increased activity. (**f**) Population-averaged early responses for drifting grating stimuli and the fitted orientation tuning curves under 5 electrical stimulation conditions. (**g**) Neurons’ early response towards pref vs. ortho gratings, under 5 electrical stimulation conditions. Red dots: mean responses. (**h**) Distributions and cumulative distributions of gOSI values of 2173 neurons under 5 electrical stimulation conditions.

To carry out population analysis, for every neuron that was visually responsive (n=2173 neurons, 3 animals, 14 FOVs), we fit its V-Stim responses with a double Gaussian tuning curve (Methods). We defined its preferred grating (denoted as “Pref” in **Fig. 2b**) as the grating, out of the eight presented to the mouse, that was closest to the peak of its tuning curve. We then defined the grating stimulus with the same orientation as the Pref grating but drifting in the opposite direction as the “Oppo” grating, which can still drive neuronal activity but not as strongly as the Pref grating. We further identified the two grating stimuli whose orientations were orthogonal to that of the Pref grating as the “Ortho” gratings, which typically evoked minimal responses from these neurons. We then averaged the ΔF/F_0_ traces for E-Stim and for Pref, Ortho, and Oppo gratings acquired under V-Stim and EV-Stim for all visually responsive L2/3 neurons (**Fig. 2c**). Ortho traces were calculated as the average of the responses evoked by the two Ortho gratings.

With E-Stim, the L2/3 population of visually responsive neurons showed early activation followed by late suppression (“No vis”, **Fig. 2c**; ΔF/F_0_ for early and late responses of individual neurons, **Supplementary Fig. 3a,b**), mirroring the same trends as in **Fig. 1f**, indicating that whether a neuron has visually evoked activity or not does not affect its response to E-Stim. With V-Stim, L2/3 population showed a steady ramp-up of ΔF/F_0_ throughout the 4 s of visual stimulation (black traces, “Pref”, “Ortho”, “Oppo”, **Fig. 2c**), with decreasing response magnitudes under Pref, Oppo, and Ortho gratings as expected.

With EV-Stim, a complex picture emerged. With ortho gratings minimally driving activity, EV-Stim with Ortho gratings closely resembled the summation of the weak E-Stim and V-Stim responses, showing early activation followed by late suppression relative to the V-Stim response (“Ortho”, **Fig. 2c**). These results indicate that with weak visually and electrically evoked activities, network activity operates in an additive regime.

For EV-stim at Pref and Oppo gratings, we observed fast-rising activity consistent with the early electrical activation, followed by a long-lasting visually evoked activity (green and purple traces, “Pref” and “Oppo”, **Fig. 2c**). However, compared to V-Stim, concurrent electrical stimulation led to a strong and sustained suppression of visually evoked activity, with the resulting activity traces clearly deviating from a simple summation of E-Stim and V-Stim responses. The suppression effect, manifested by the decrease in ΔF/F_0_, was greater at higher electrical currents and with stronger visually evoked activity (e.g., larger ΔF/F_0_ decrease in “Pref” than “Oppo”, **Fig. 2c**), suggestive of a divisive inhibitory mechanism^40,41^ at work.

### Electrical stimulation reduces orientation selectivity of L2/3 V1 PYR neurons

Because electrical stimulation increased the activity of L2/3 neurons during non-preferred visual stimulation and decreased the activity during preferred visual stimulation, we expected that combining electrical and visual stimulation would reduce the orientation selectivity of L2/3 neurons.

We sorted all visually responsive L2/3 neurons by their preferred drifting directions under V-Stim and plotted their early responses for all grating stimuli (**Fig. 2d**). Without electrical stimulation, the plot revealed a diagonal line of peak early response ΔF/F_0_ (red bars in middle column “0”, **Fig. 2d**), indicating that L2/3 neurons are mostly orientation-tuned with their preferred orientation/direction evenly distributed, in agreement with previous studies^33,42^.

With concurrent electrical stimulation, the diagonal line became more difficult to visualize (“200_C_”, “100_C_”, “100_A_”, “200_A_” columns, **Fig. 2d**), due to both the suppression of the early response to the preferred grating and the increase in activity at the non-preferred gratings. This trend became even more apparent when we subtracted the V-Stim responses from the EV-Stim responses, where blue diagonal lines indicated the suppression of visually evoked activity at the preferred grating by electrical stimulation (**Fig. 2e**). This suppression lasted throughout the duration of visual stimulation, persisting for several seconds after the end of the 10-ms-long electrical stimulation, as indicated by the late responses of individual neurons (**Supplementary Fig. 3c, d**).

The population tuning curves, which were double-Gaussian fits to the early responses of all neurons showed the same trends: electrical stimulation increased the activity at the non-preferred orientations (“Ortho” and “Oppo”, **Fig. 2f**) while decreasing the activity at preferred orientation (“Pref”, **Fig. 2f**) (p values, **Supplementary Table 1**). Therefore, as a population, L2/3 neurons had reduced orientation selectivity that became more severe at higher electrical stimulation currents.

The reduction of orientation selectivity can also be appreciated in the scatter plots of the early responses to Pref versus Ortho gratings for individual neurons (**Fig. 2g**). Electrical stimulation caused the response distributions to shift toward a decrease in Pref response and an increase in Ortho response. Consistently, we observed a significant decrease in the global orientation selectivity index (gOSI) distributions that became more pronounced at higher stimulation currents (**Fig. 2h**; directional Wilcoxon ranksum test; p values, **Supplementary Table 3**).

In these experiments, L2/3 neurons, whether excitatory or inhibitory, were indiscriminately labeled with GCaMP6s. However, the population response should be dominated by excitatory PYR neurons due to their comprising the vast majority of L2/3 neurons^43,44^. Indeed, when we applied additional criteria to restrict our analysis to putative PYR neurons (i.e., skewness > 2.7^45^ or gOSI > 0.3^33^), similar results were obtained (**Supplementary Fig. 4**).

Together, our measurements indicate that with or without simultaneous visual stimulation, electrical stimulation transiently and rapidly activates L2/3 PYR neurons followed by a long-lasting inhibition. When L2/3 PYR neurons transition from low firing rates to high firing rates in response to visual stimulation, the sustained inhibition by electrical stimulation transitions from a subtractive to a divisive regime. The early direct electrical activation, together with the stronger inhibition that occurs when the neurons respond to their preferred visual stimuli, leads to a reduction in orientation selectivity.

### PV interneurons are not excited by electrical stimulation alone and their visually evoked activity is suppressed by electrical stimulation

Whereas the rapid and direct activation of L2/3 PYR neurons by electrical stimulation can be explained by the activation of their axons in L1 which are in close proximity to the stimulating electrode, the long-lasting inhibition following electrical stimulation requires the involvement of INs. Given that different IN subtypes have distinct roles in regulating cortical activity, we labeled distinct IN populations by using Cre-dependent expression of GCaMP6s in V1 of subtype-specific Cre transgenic lines and carried out similar experiments.

We first investigated Parvalbumin-expressing (PV) INs, which play major roles in cortical inhibition by innervating the cell bodies and basal dendrites of nearby PYR neurons^44^ and are known to mediate divisive inhibition^41^. We injected AAV2/1-Flex-Syn-GCaMP6s into transgenic Pvalb-IRES-Cre^34,35^ mice to selectively express GCaMP6s in L2/3 and L4 (300 – 420 µm below dura) PV INs and imaged their calcium activity under E-Stim, V-Stim, and EV-Stim conditions (3 animals, 6 FOVs, 276 L2/3 neurons, 381 L4 neurons; example FOV, **Fig. 3a**).

**Figure 3.**
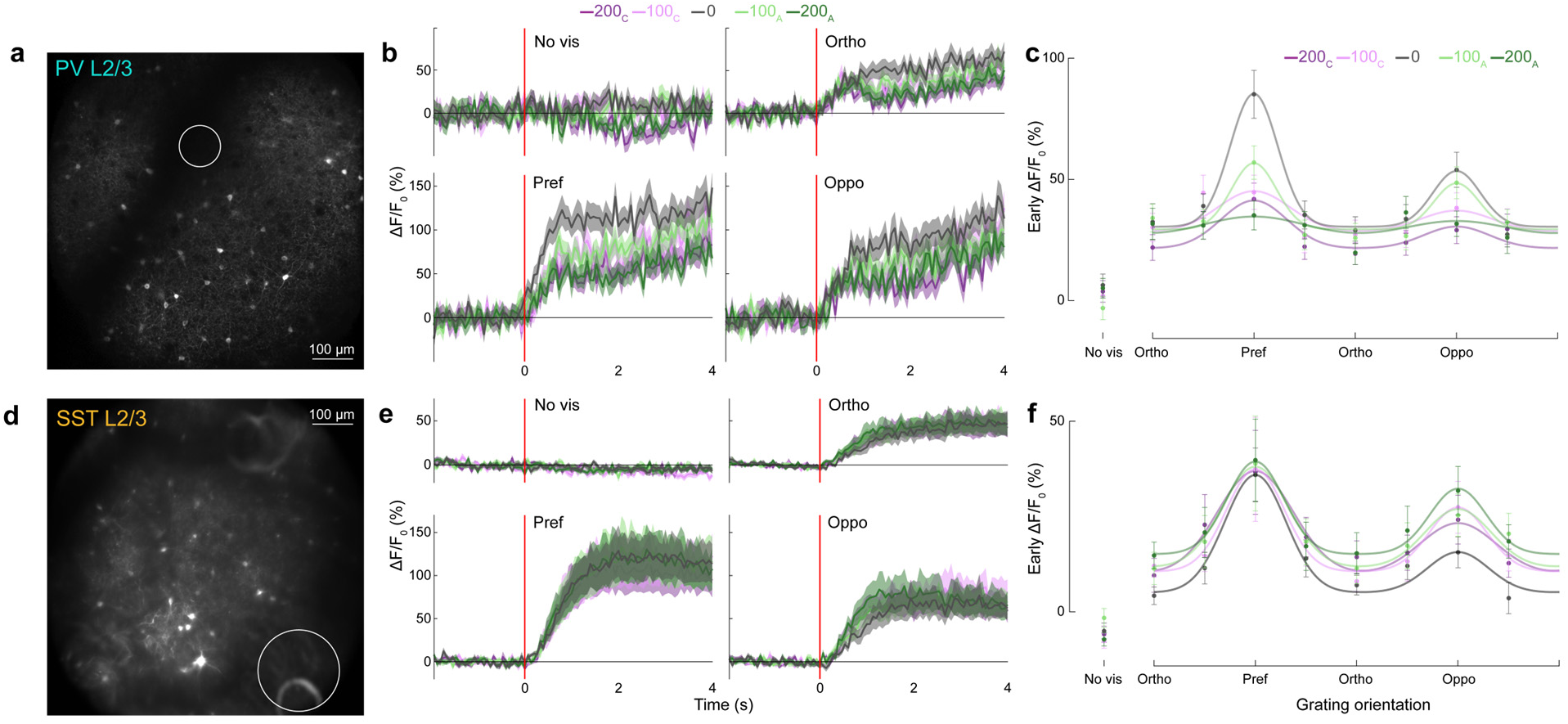
– Electrical stimulation suppresses visually evoked activity in PV interneurons and weakly enhances the visually evoked activity in SST interneurons. (**a**) Example 2PFM image of PV INs expressing GCaMP6s 180 µm below electrode surface in a PV-ires-Cre mouse. Circle: profile of stimulation electrode. (**b**) Population- and trial-averaged ΔF/F_0_ traces of 156 L2/3 PV INs with visually-evoked activity under 5 electrical stimulation conditions, grouped for 4 visual stimulation conditions: no stimulus, ortho, pref, and oppo. Trace and shade: mean and SEM. Red lines: stimulus onset. (**c**) Scattered data points: Early ΔF/F_0_ of the traces in **b**; tuning curves: double-Gaussian fits to the scattered data. (**d-f**) Same as **a-c**, but for SST INs with visually-evoked activity. **d**: example image acquired at 160-230 µm below electrode surface in an SST-ires-Cre mouse; **e-f**: analysis on 274 SST INs with visually evoked activity.

We averaged the ΔF/F_0_ traces of 156 visually responsive L2/3 PV INs (**Fig. 3b**). Interestingly, E-Stim did not evoke a calcium response from PV INs (“No vis” panel, **Fig. 3b**), likely because neither the PV IN cell bodies nor their neuronal processes (and especially their axons) were sufficiently close to the cortical surface electrode to be excited. However, L2/3 PV INs are known to receive strong recurrent excitation from L2/3 PYR. The lack of PV IN activation by E-Stim therefore suggests that there is an additional short latency inhibitory input preventing them from firing (see NDNF section below). V-Stim evoked dynamics in PV INs similar to those observed in L2/3 neurons, with a steady increase in ΔF/F_0_ throughout the 4 s of visual stimulation and decreasing response magnitudes under Pref, Oppo, and Ortho gratings (black traces, “Pref”, “Ortho”, “Oppo” panels, **Fig. 3b**).

When combined with visual stimulation, electrical stimulation suppressed visually evoked responses of PV INs to Pref, Oppo, and Ortho gratings (green and purple traces, “Pref”, “Ortho”, “Oppo” panels, **Fig. 3b**), with stronger inhibition observed at larger electrical currents (**Fig. 3c**). Therefore, electrical stimulation inhibits PV neuron activity regardless of the strength of their concurrent sensory-evoked responses. This is different from L2/3 PYR neurons, for which electric stimulation suppresses the activity at the preferred orientation but increases the activity at non-preferred orientations. The same trends were observed in individual neurons (**Supplementary Fig. 5a-c**). The suppression of PV activity by electrical stimulation suggests that PV INs are unlikely to be the source of the late-onset inhibition observed in L2/3 PYR neurons.

### SST interneurons are not excited by electrical stimulation alone and their visually evoked activity is weakly enhanced by electrical stimulation

We next investigated whether the inhibition of PYR neuron activity may instead be mediated by somatostatin (SST)-expressing L2/3 INs. Being Martinotti cells, these SST INs are one of the main inhibitory inputs to PYR apical dendrites through their axonal projections to L1^44^. These projections were close to our electrode, potentially enabling the antidromic activation of SST neurons by the ECoG electrode.

To test this hypothesis, we injected AAV2/1-Flex-Syn-GCaMP6s into transgenic SST-IRES-Cre mice^36^ to selectively express GCaMP6s in SST INs within L2/3 and L4 and measured their responses to E-Stim, V-Stim, and EV-Stim (5 animals, 21 FOVs, 368 L2/3 neurons, 470 L4 neurons; example FOV, **Fig. 3d**). Due to the sparsity of SST IN cell bodies in V1, we used Bessel-focus 2PFM, a fast volumetric imaging method^46^, to improve imaging throughput (Methods). We focused further analysis on L2/3 SST INs with visually evoked activity (274 neurons).

Surprisingly, we did not observe calcium transients caused by direct activation of L2/3 SST INs under E-Stim (“No vis” panel, **Fig. 3e**). Under V-Stim, SST INs exhibited similar temporal activity dynamics as PV INs but had their ΔF/F_0_ saturate in magnitude ∼2 s after visual stimulation onset (black traces, “Pref”, “Ortho”, “Oppo” panels, **Fig. 3e**).

During EV-Stim of visually responsive L2/3 SST INs, electrical stimulation either enhanced or suppressed a neuron’s early response to its preferred stimuli (**Supplementary Fig. 5d,e**). Averaging the ΔF/F_0_ traces of all visually responsive L2/3 SST INs, we observed weak and statistically insignificant ΔF/F_0_ increases in the population activity traces at preferred gratings, and small but statistically significant ΔF/F_0_ increases for non-preferred gratings relative to V-Stim traces (**Fig. 3e**; Early response, **Fig. 3f**, **Supplementary Fig. 5f**, **Supplementary Table 1**).

One explanation for the observed weak electrical activation during visual stimulus presentation could be that axon segments of SST INs were directly activated by electrical stimulation but the antidromic activation was too weak to induce firing at the SST somata. However, with concurrent excitatory inputs from visual stimulation, subthreshold activation by electrical stimulation could lead to an increase of SST IN firing rates. This increase was more pronounced for the non-preferred gratings, as the weaker sensory drive under these conditions enabled the subthreshold activation by electrical stimulation to make a more appreciable impact on firing rates.

We observed similar trends for the E-Stim, V-Stim, and EV-Stim responses of L4 PV and SST INs (**Supplementary Fig. 6**): Neither PV nor SST INs were activated by electrical stimulation alone; when combined with visual stimulation, electrical stimulation suppressed the visually evoked activity of PV INs across all visual input strengths and moderately increased the activity of SST INs only under weaker visual input at non-preferred gratings. Therefore, neither IN subtypes can mediate the slow-onset, long-lasting inhibition observed under E-Stim and EV-Stim.

### Electrical stimulation first activates and then inhibits NDNF IN activity

Another inhibitory cell type that targets PYR apical dendrites is L1 NDNF INs^47,48^. We injected AAV2/1-Flex-Syn-GCaMP6s into transgenic NDNF-IRES-Cre mice to selectively express GCaMP6s in INs expressing neuron-derived neurotrophic factor (NDNF) protein, which represent ∼70% of L1 IN population^37^, and measured their responses to E-Stim, V-Stim, and EV-Stim (3 animals, 15 FOVs, 884 neurons; example FOV, **Fig. 4a**).

**Figure 4.**
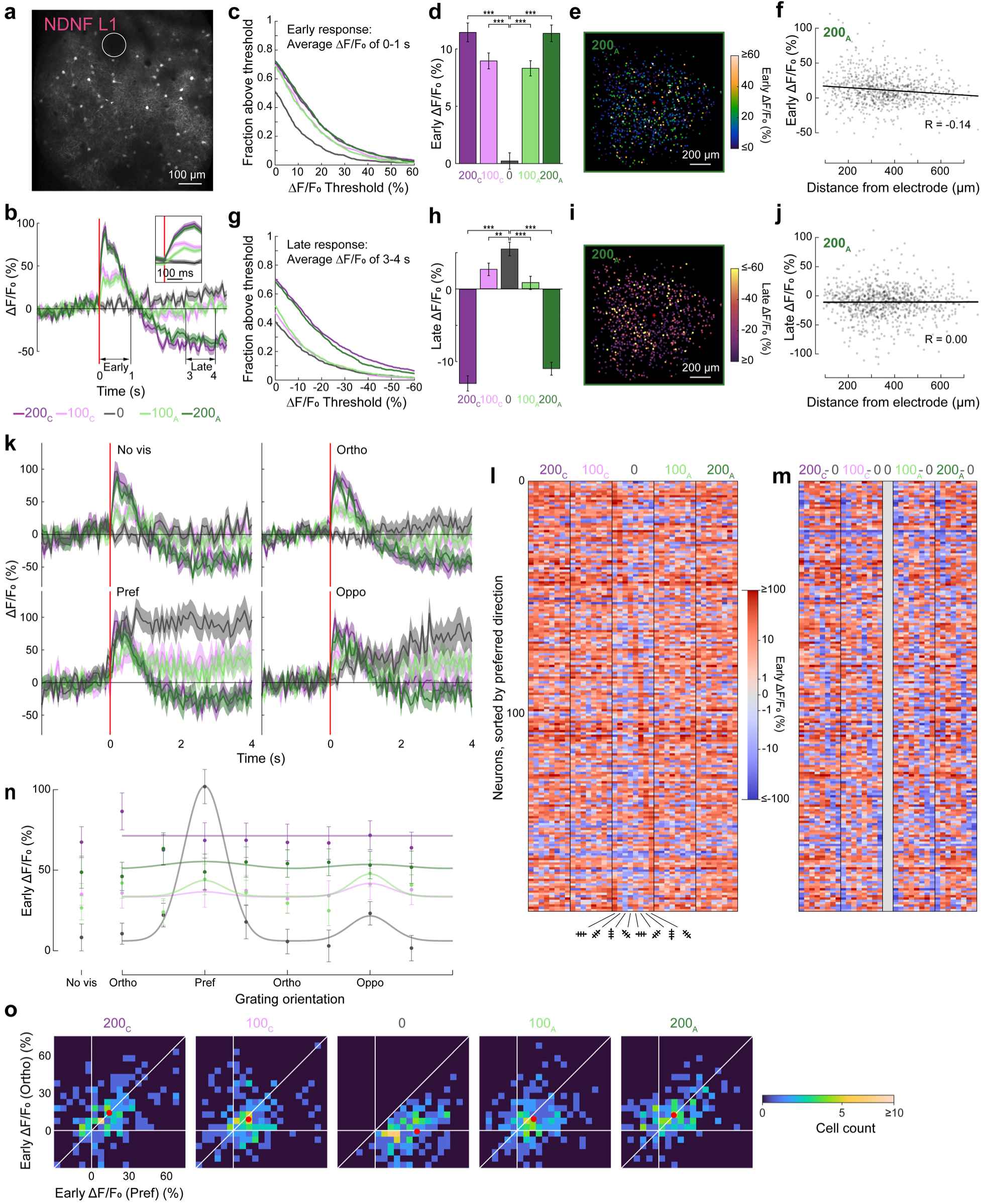
Cortical-surface electrical stimulation strongly modulates activity and orientation selectivity of L1 NDNF INs. (**a**) Example 2PFM image of NDNF INs expressing GCaMP6s 90 µm below electrode surface in a NDNF-ires-Cre mouse. Circle: profile of stimulation electrode. (**b**) Average ΔF/F_0_ traces of all L1 NDNF neurons (N = 884) in response to E-Stim (red line). Trace and shade: mean and SEM. Inset: zoomed-in to 0-200 ms post E-Stim. (**c**) Fractions of neurons with their early response (average ΔF/F_0_ during 0-1 s post E-Stim) above a specific threshold value vs. threshold values, for all stimulation currents. (**d**) Time-averaged early response ΔF/F_0_ of 884 neurons. Error bar: SEM. Two-sample KS-test; P values: **: < 0.01; ***: < 0.001. (**e**) Early response to 200_A_ E-Stim, plotted for each neuron based on their lateral displacement from electrode (red “+”: electrode center). (**f**) Early response to 200_A_ E-Stim of each neuron vs. its 3D distance from electrode center. R: Pearson’s correlation coefficient. (**g,h,i,j**) Same as (**c,d,e,f**) but for time-averaged late response ΔF/F_0_. (**k**) Population- and trial-averaged ΔF/F_0_ traces of 185 L1 NDNF neurons with visually evoked activity under 5 electrical stimulation conditions, grouped for 4 visual stimulation conditions: No visual stimulus, Ortho, Pref, and Oppo. Trace and shade: mean and SEM. Red lines: stimulus onset. (**l**) Heatmap of early response ΔF/F_0_. Rows: individual cells sorted by their preferred direction; Columns: electrical-visual stimulation conditions. (**m**) Heatmap of the difference in early response ΔF/F_0_, by subtracting V-Stim responses from EV-Stim responses in **j**. Blue: reduction in response; red: increase in response. (**n**) Scattered data points: early-response ΔF/F_0_ of the traces in **k**; tuning curves: double-Gaussian fits to the scattered data. (**o**) Neurons’ early response towards Pref vs. Ortho gratings, under 5 electrical stimulation conditions. Red dots: mean responses.

Mirroring the activity dynamics of L2/3 PYR neurons but with much larger ΔF/F_0_ magnitudes, NDNF INs showed immediate and strong activation by E-Stim (**Fig. 4b**, 884 neurons) with calcium activity rising rapidly within 100 – 200 ms (e.g., reaching 100% for 200 µA stimuli), followed by a strong inhibition that reduced activity below baseline (e.g., to −50% ΔF/F_0_ for 200 µA stimuli). Both 100 µA and 200 µA stimulation currents strongly activated neurons, as indicated by the high fractions of neurons with their average ΔF/F_0_ during 0-1 s post E-Stim above threshold values (**Fig. 4c**). Early responses of NDNF INs to E-Stim were significantly greater than those without stimulation (two-sample KS-test, **Supplementary Table 1**) and responses to 200 µA anode-first stimulation currents significantly higher than those to 100 µA anode-first stimulus (two-sample KS test; p value: 200_C_ vs. 100_C_: 5.6×10^−2^; 200_A_ vs. 100_A_: 7.7×10^−4^, **Fig. 4d**). Unlike PYR neurons, there were no significant differences between early responses to cathode-leading and anode-leading E-Stim (**Fig. 4d**, **Supplementary Table 1**). We observed a weak negative correlation between NDNF INs’ early response strength and their distance from the center of the stimulating electrode for all electrical stimulation conditions (200_A_ data, **Fig. 4e,f**, two-tailed Pearson’s correlation significance test, p = 4.4×10^−5^; Other conditions, **Supplementary Fig. 7a**). The late-onset inhibition of NDNF INs induced by E-Stim was strongest at 200_A_ and 200_C_ (**Fig. 4g,h**) and, similar to the late-onset inhibition of PYR neurons, showed no significant correlation with distance from the electrode center (200_A_, **Fig. 4i,j**; Other conditions, **Supplementary Fig. 7b**).

### Electrical stimulation reduces orientation selectivity of NDNF INs and leads to late-onset long-lasting inhibition of their visually evoked activity

When excited by V-Stim, NDNF INs with visually evoked activity exhibited strong direction preference, consistent with previous findings^49^ (N = 185; population response, black traces, **Fig. 4k**; individual neuron early response, middle column, “0”, **Fig. 4l**). At their preferred grating stimuli, NDNF INs had sustained visually evoked responses throughout the 4-s duration of visual stimulation (black trace, “Pref” panel, **Fig. 4k**).

Compared with V-Stim responses, when electrical stimulation of 100 µA currents was combined with visual stimulation, NDNF INs had suppressed early responses at Pref gratings but elevated early activity at all non-preferred gratings (population response, light green and light purple traces, **Fig. 4k**; individual neuron response, **Fig. 4l,m**). At 200 µA currents, electrical stimulation dominated visual stimulation, leading to ΔF/F_0_ responses that resembled those evoked by electrical stimulation alone for both preferred and non-preferred stimuli (cf. population response, dark green and dark purple traces in “Pref”, “Ortho”, “Oppo” panels to those in “No vis” panel, **Fig. 4k**; individual neuron response, **Fig. 4l,m**).

As a result, the population tuning curves fitted to the early-response ΔF/F_0_ for NDNF INs showed that electrical stimulation led to an almost complete loss of orientation/direction tuning (**Fig. 4n**). The scatter plots of the early responses to Pref and Ortho gratings for individual neurons showed a reduction in Pref grating response and an increase in Ortho grating response, causing the response distributions to shift towards the diagonal line representing equal responses (**Fig. 4o**).

### Circuit model for cortical electrical stimulation and its interaction with concurrent sensory processing

In order to explain the above observations in a manner consistent with the anatomy and physiology of mouse V1, we propose a simplified circuit model of how electrical stimulation affects the activity of cortical neurons and alters how they respond to concurrent visual stimulation (**Fig. 5**).

**Figure 5.**
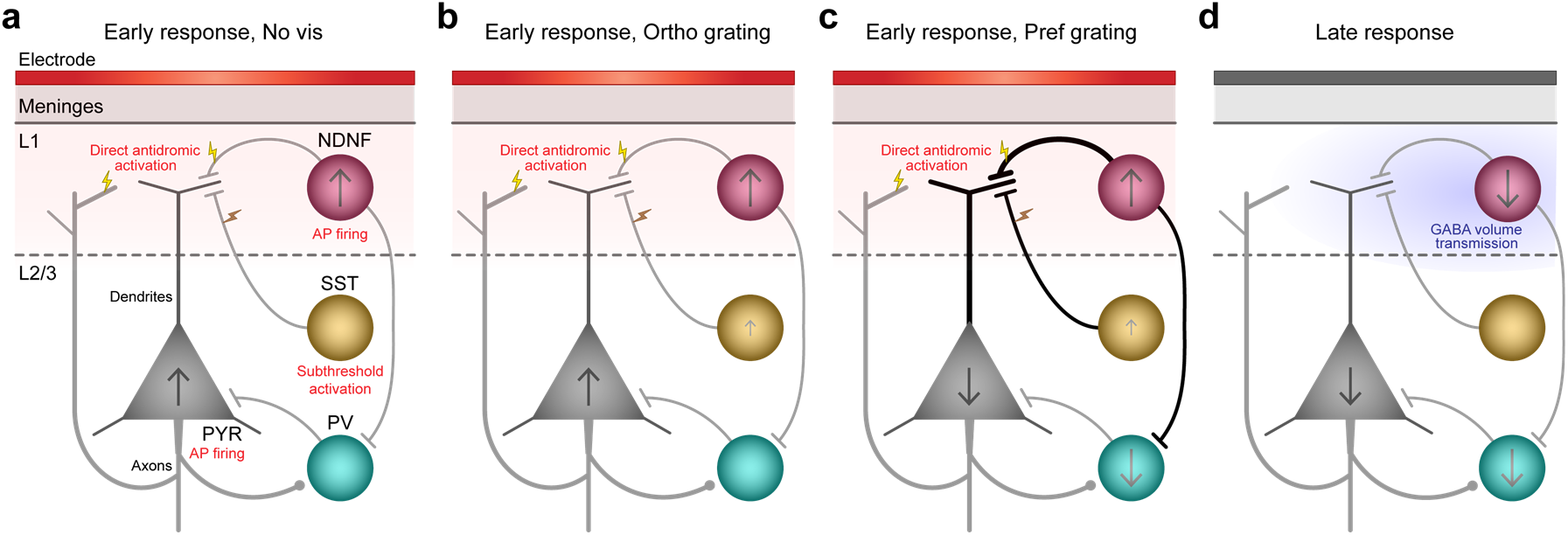
Proposed circuit model during cortical electrical stimulation and concurrent visual stimulation. Electrode is above mouse V1. Gray, cyan, yellow, and magenta objects represent PYR neurons, PV, SST, and NDNF INs. Arrows within cells represent change in firing rate. Light red shade: direct electrical stimulation’s region of influence; Yellow lightning bolts: direct antidromic activation; Orange lightning bolts: direct subthreshold activation. (**a**) Early response to electrical stimulation, in the absence of visual stimulation. (**b**) Early response to electrical stimulation, under weak visual stimulation (e.g., evoked by Ortho grating). (**c**) Early response to electrical stimulation, under strong visual stimulation (e.g., evoked by Pref grating). (**d**) Late suppression response mediated by GABA volume transmission (light blue shade) released by early NDNF activation.

In this model, cortical surface electrical stimulation by itself activates PYR neurons and NDNF INs mainly through direct antidromic activation of their axon segments in L1, leading to AP firing and a rapid increase in their intracellular calcium during the first second after electrical stimulation. It also leads to subthreshold activation of SST INs but does not activate PV INs (**Fig. 5a**). (Even though only antidromic activation is considered in this model, other activation mechanisms can be also engaged – see Discussion.)

When concurrent visual stimuli are at the orthogonal, non-preferred orientations of PYR neurons and NDNF INs, these neurons are minimally excited by visual stimulation. Their early-stage calcium dynamics are therefore similar to those observed under electrical stimulation alone (**Fig. 5b**). Because SST INs are less orientation tuned than PYRs and NDNF INs^49^, they have comparatively stronger visually evoked activity at Ortho gratings, which adds up with their subthreshold activation by electrical stimulation, leading to a moderate increase in firing rate.

When visual stimuli are presented at the preferred direction of PYRs and thus strongly activate them, their otherwise strong visually-driven activity is suppressed by the electrically activated NDNF INs, which reduce PYR activity by inhibiting their apical dendrite directly (**Fig. 5c**)^47,48^ (and by blocking excitatory inputs on PYR via presynaptic GABA_B_ receptors; see **Fig. 5d**). The suppression of PV activity arises from both the reduction in PYR activity, which provides less driving force for PV neuron firing, and mainly extrasynaptic inhibition by NDNF INs^49–51^.

Once activated, NDNF INs release GABA into the extracellular space via volume transmission^52^ and lead to sustained inhibition (**Fig. 5d**), likely through GABA_B_ receptor-mediated responses in their target cell types including PYR neurons, PV INs, and NDNF INs themselves. Such suppression of activity persists for seconds after electrical stimulation and is stronger at larger stimulation currents.

In summary, this model suggests that early activation of PYR and NDNF neurons was principally due to direct activation of L1 axons near the stimulation electrode, while the later prolonged inhibition results from volumetric GABA cloud released by the activated NDNF INs. Overall, our results are consistent with electrical stimulation engaging the top-down modulatory functions of L1 NDNF INs, which have been extensively documented in the recent literature^53^.

## Discussion

Using in vivo calcium imaging of the awake mouse brain with chronically implanted ECoG electrodes, we measured how electrical stimulation modulates the activity of visual cortical neurons, including during concurrent visual stimulation. We found that cortical surface electrical stimulation first activates L2/3 neurons, followed by a prolonged inhibition that lasts seconds after stimulation. Electrical stimulation suppresses the activity of PYR neurons at their preferred grating orientation but increases their activity during non-preferred visual stimulation, thereby reducing sensory feature selectivity. Among inhibitory subtypes, we found that electrical stimulation does not activate PV INs, activates SST INs in a subthreshold manner, and strongly activates NDNF INs. We propose a circuit model in which L1 NDNF INs are strongly activated by cortical electrical stimulation and subsequently inhibit L2/3 PYR neurons and PV INs, likely through volume transmission of GABA.

### Fast-onset direct activation by electrical stimulation

Our in vivo calcium imaging experiments indicate that, under our stimulation protocol, cortical-surface electrical stimulation directly activates L2/3 PYR neurons and L1 NDNF inhibitory neurons, leading to action potential firing and an increase in their intracellular calcium concentration. It also leads to subthreshold depolarization of L2/3 and L4 SST INs, with the activation only measurable when combined with visual stimulation. In contrast, electrical stimulation does not activate L2/3 or L4 PV INs.

These observations are consistent with biophysical studies that found axon initial segments or nodes of Ranvier have the lowest activation threshold to external electrical input^22–24^. With the ECoG electrodes placed on cortical surfaces, neurons with axonal projections in L1 can therefore be directly activated electrically. (L1 NDNF neurons could also be activated synaptically if stimulation induced orthodromic action potentials in L1 horizontal fibers; see next section).

Indeed, we found that NDNF neurons, with dense horizontally extending axonal arbors in L1^37^, had the largest ΔF/F_0_ amplitude in response to electrical stimulation in our study, suggesting strong activation by the cortical-surface electrode. The smaller ΔF/F_0_ amplitudes evoked by electrical stimulation in L2/3 PYR neurons indicate that they were more moderately activated, likely because their local axon projections, although present in L1, are mostly concentrated in lower L1 and upper L2/3^54^. For L2/3 PYR neurons and L1 NDNF INs, we observed more neurons activated at larger stimulation currents and stronger activation in neurons with cell bodies closer to the stimulation electrode, similar to previous experimental observations with intracortical electrical stimulation^27,28,55,56^ and consistent with simulations based on biophysical models^57^.

L2/3 and L4 SST-expressing Martinotti cells have ascending axons into L1^44,58–60^. Our data suggests that electrical stimulation employed here leads to subthreshold depolarization at the SST IN cell bodies, which does not lead to detectable calcium transients by itself but can increase firing rate when combined with sensory stimulation. In contrast, the axon projections of L2/3 and L4 PV INs are mostly within L2/3 and L4^58,60^, likely too far away from the cortical surface to be activated under our stimulation protocol. This explains the lack of direct activation of PV INs by electrical stimulation.

Comparing stimuli with the same current but opposite polarities, we observed a small but significant increase in ΔF/F_0_ of PYR neurons in response to cathode-leading stimuli. This is consistent with previous findings in animal models^55,56,61^, as well as clinical observations of cathode-leading stimulation being associated with greater treatment efficacy^62,63^. The smaller difference between cathode and anode-leading pulses observed here was likely due to the fact that by placing the ECoG electrode on cortical surface, the stimulation current vectors interacted with axons at a large variety of orientations, which minimized the effects of polarity. In contrast, previous studies stimulated through an electrode tip embedded in the brain tissue and observed stronger polarity effects in the immediate vicinity of the electrode, where the orientation of the current vector relative to the membrane of excitable elements was fixed. Another difference could be our use of a relatively long stimulation pulse wherein the second phase greatly exceeds the refractory period from the first phase so effects of both polarities should be produced. Finally, we used a single stimulation pulse instead of a train stimulus. The cumulative effects from a train of pulses might amplify response differences and lead to larger differences between polarities.

### Other activation mechanisms by electrical stimulation for L1 NDNF INs

Our simplified circuit model above only considered *antidromic axonal* activation, in which activation of NDNF axons leads to direct action potential invasion of the soma. Given that NDNF INs have dense axons and dendrites within L1^37^ in close proximity to the ECoG stimulation electrode, they may also be activated via other mechanisms^30^ including *orthodromic axonal* with synaptic activation, where afferent axons carrying synaptic input to NDNF cells were activated by electrical stimulation, and *direct depolarization* of the dendritic/somatic membrane by the impressed electrical currents. Several characteristics of NDNF activation support the involvement of orthodromic activation.

Our data indicate that antidromic invasion of the NDNF somata was incomplete. Anatomical studies show that L1 NDNF cells have a very dense axonal arbor within 200 µm from the soma^64^. The lack of response by many NDNF cells less than 200 µm from the electrode (**Fig. 4f**, **Supplementary Figure 7a**), even at high stimulation currents, suggests that antidromic spikes are evoked but fail to invade the soma.

A cell responding to pure antidromic stimulation at a lower amperage (e.g., 100 μA) would not increase its firing rate at higher amperages (e.g., 200 μA). This is because even if multiple axon branches were stimulated at higher amperages, antidromic stimulation would not result in multiple action potentials at the axon hillock, as the final common path would be refractory when the slightly later conducting ones arrive^65^. However, we observed larger ΔF/F_0_ for 200 µA stimuli than for 100 µA stimuli (**Fig. 4c**), consistent with orthodromic activation with amperage-dependent responses.

L1 NDNF neurons in V1 receive excitatory input mainly from thalamic and other cortical areas^49,53^, and inhibitory input from SST (Martinotti) cells as well as other L1 NDNF cells^51,66^. Both could be activated orthodromically by electrical stimulation, evoking or inhibiting cell firing directly. These orthodromic inputs could also interfere with invasion of the antidromic spikes. The afferent thalamic and cortical axons have a higher level of myelination and thus lower threshold for electrical activation than the axons of NDNF and SST cells. (Although the ascending segment of the SST axons innervating NDNF neurons are myelinated, their arborization in L1 is not^66^.) Therefore, the net effect of orthodromic activation is excitatory, as we observed in the early response of NDNF INs.

Direct depolarization of the neuronal membrane seems to be ruled out by our data. For dendrites at the same distance from the stimulating electrode, depolarization due to direct activation should have similar magnitude and thus similar likelihood of activation. Given the decrement of the tissue current vectors and that the dendrites of NDNF cells are nearly all <100µm from the soma^64^, direct dendritic activation should also drop very quickly with distance. However, we observed spotty distribution and a slow decay of activated cells with distance (**Fig. 4e,f**).

### Reduction of visual feature selectivity by electrical stimulation

Stimulation through ECoG electrodes in sensory cortex is known to generate sensory perception in humans^9–11,67,68^. In the awake mouse, the direct activation of V1 PYR neurons by electrical stimulation may similarly lead to visual perception. However, previous studies showed that when combined with visual stimulation, intracortical electrical stimulation in visual cortical areas can impair performance in visual discrimination^69–71^ and memory^72,73^. This is consistent with our observation that electrical stimulation applied at the onset of visual stimulation reduces selectivity for drifting grating orientations and directions for all cell types investigated.

For L2/3 PYR neurons and L1 NDNF INs, electrical stimulation suppresses their population activity during preferred grating stimulation and increases their activity during non-preferred grating stimulation, resulting in more similar firing rates for distinct grating stimuli and a reduction of their gOSIs.

Electrical stimulation suppresses the firing of PV INs and leads to subthreshold activation of SST INs. Previously, it was reported that activating V1 PV and SST INs improves feature selectivity and visual perception in awake mice^74,75^, (but see conflicting results and discussions^41,76–78^). Given the strong reduction of orientation tuning observed in PYR neurons, we speculate that electrical stimulation reduces the mouse’s orientation discrimination ability, a speculation that should be further tested through behavior assays^74,77,79,80^.

### L1 NDNF neurons mediate inhibition by electrical stimulation

For L2/3 PYR neurons, electrical stimulation impacts neuronal activity in two different regimes. For PYR neurons that are not or only weakly activated by visual stimulation, direct electrically evoked response adds to the visual response. When the neurons are strongly activated by visual stimulation, concurrent electrical stimulation leads to a decrease in firing rates in a manner that is consistent with divisive inhibition, which requires the involvement of INs within the local circuit.

By selectively measuring the responses of IN subtypes with calcium imaging, we identified L1 NDNF INs as the main mediator of this selective inhibition. In V1, NDNF cells receive bottom-up visual information (e.g., from cells within V1 and the dorsal lateral geniculate nucleus) and top-down inputs (e.g. from other sensory, motor, association, and prefrontal cortices, higher-order thalamic nuclei, and hypothalamus) from multiple brain areas^49,81^. Representing ∼70% of L1 INs, NDNF-expressing cells consist of neurogliaform cells and canopy cells^37,47,82^. These cells have dense L1 axonal projections that extend horizontally, through which they interact with nearby PYR neurons and other IN subtypes and act as the master top-down regulator of cortical circuits^83^.

As discussed above, NDNF INs can be activated by cortical surface electrodes through both antidromic and orthodromic mechanisms. Previous studies indicate that NDNF activation leads to divisive inhibition of PYR neurons through their dendritic tufts, as well as the inhibition of PV but not SST INs^47–49,83^, effects that are consistent with our experimental observations here.

The long-lasting inhibition observed here was likely mediated by the neurogliaform cells within the NDNF population. GABA released from their dense axonal arborizations can elicit inhibitory responses from nearby neurons both through classical synaptic transmission and extrasynaptically through GABA released into the extracellular space as a volume transmitter^52,84^. Uniquely among GABAergic INs, even the firing of a single action potential by neurogliaform cells can produce slow inhibition in their target neurons through GABA_B_ receptors along the dendritic arbors of PYR neurons and INs^44^.

Given the large ΔF/F_0_ transients evoked by E-Stim from NDNF INs, it is reasonable to speculate that a single cortical-surface electrical stimulation pulse can strongly activate and potentially trigger persistent firing from neurogliaform INs^83^, resulting in the observed strong and long-lasting inhibition of nearby neurons, including NDNF INs themselves, through volume transmission of GABA^85^. Here, the involvement of volume transmission, as well as the fact that their axons extend hundreds of microns horizontally, explains the lack of distance dependence of the late-onset inhibition. The involvement of GABA_B_ receptors and volume transmission will be tested pharmacologically in future work.

Whereas optogenetic activation of L1 NDNF INs leads to hyperpolarization that lasts several hundred milliseconds in excitatory neurons across cortical layers 2-5^49^, the inhibitory effects induced by electrical stimulation last several seconds, suggesting cortical-surface electrical stimulation as a powerful switch capable of decreasing or even muting outputs from an entire cortical column^86^. Through presynaptic GABA receptors on both locally projecting and long-range axons^87–92^ in L1, the long-lasting inhibition following the early-onset excitation, induced by volume transmission of GABA, could also modulate feedforward, feedback, and neuromodulatory inputs carried by these projections to V1.

Intriguingly, L1 INs in the human and mouse neocortex have similar physiology, subtypes, marker expression (i.e., highly enriched NDNF-expressing cells in both human and mouse L1), and responses to neuromodulation^93^, suggesting that L1 INs in the human neocortex may carry out comparable functions. Whether cortical electrical stimulation similarly modulates human cortical activity requires further investigation^94,95^.

Finally, while the local effect of cortical electrical stimulation is dominated by inhibition, its impact on global activity may be more varied. L1 contains massive projections from a diverse set of cortical and subcortical brain regions, including those involved in neuromodulation. Through antidromic activation of these L1 axons, electrical stimulation could modulate the firing rates and excitability of their cell bodies, leading to brain-wide changes in neural dynamics, which may be evaluated with mesoscale measurements in the future.

### Outlook and future work

In this study, we investigated how a single 10-ms-long symmetric biphasic pulse, either anode- or cathode-leading with 100 µA or 200 µA current, modulates the activity of cortical neurons, including during visual stimulation. Our experimental approach – imaging neuronal activity with cell-type specificity from awake mice with chronically implanted ECoG electrode during electrical and sensory stimulation – can be used to explore the parameter space of stimulation protocols (e.g., frequency, polarity, and waveform) for generating cortical activation/inhibition with desired spatial and temporal patterns.

The above in vivo imaging experiments are synergistic with computational approaches that have been developed to predict neuronal responses to electrical stimulation from cortical surfaces, ranging from biophysics simulations of individual anatomically accurate neurons^57,96,97^ to models incorporating cortical networks^57,98^. These approaches predict how different neuronal subtypes respond to the same electrical stimulus, through direct activation and/or indirectly through modulation of circuit activity. The existing modeling efforts have focused on NHPs^97,98^ and rats^57,96^, but not mice, which are likely the best model system for studying electrical stimulation of the brain due to their genetic toolkit and optical accessibility. An ideal computational model developed for the mouse cortex would account for the complete cortical circuit, including all known major neuronal subtypes in all layers (but especially L1 NDNF INs). It should also include representations of bottom-up sensory input and top-down modulation associated with brain states and ongoing activity. Models that succeed in predicting activity patterns in the mouse cortex may then be applied to human cortex using information on human cortical cell types, morphologies, and circuit connectivity. Similarly, to utilize electrical stimulation for therapy in awake and behaving patients, models should also consider interactions between electrically evoked and sensory evoked activity, as well as incorporate behavioral states (e.g., arousal levels) and/or ongoing activity, which are known to modulate sensory-evoked activity^99^ and electrically evoked activity^29^ in the mouse brain.

Finally, our study highlights the value of 2PFM as a tool for measuring neuronal response to electrical stimulation. Our results set the stage for future research efforts, including validating models of neuronal responses to electrical stimulation^57,96–98^, as well as designing stimulation protocols for spatially and temporally varying activation and inhibition of specific cell type/circuits. The advent of population voltage imaging capability^100^ will further enable studying electrical-stimulation-evoked activity at millisecond time resolution and subthreshold sensitivity. Combined with measurements of neurotransmitter and neuromodulator release in response to electrical stimulation using genetically encoded fluorescent sensors, 2PFM is poised to continually contribute to our circuit and molecular understanding of the effects of electrical stimulation.

## Author Contribution

N.J. and E.H. conceived of the project; J.L.F, N.J., and E.H. designed the experiments; J.L.F. performed surgery, acquired and analyzed all imaging data; K.L., Y.T., M.G., R.V., and S.D. fabricated and tested the electrodes; J.G. developed the stimulation protocols; J.L.F. and H.Y.Y. measured ECoG electrode properties; J.L.F., N.J. and E.H. wrote the manuscript with input from all authors.

## Acknowledgement

We thank Anna Devor lab for helpful suggestions on implanting ECoG electrodes. This work was supported by NIH BRAIN® Initiative R01NS109553 (N.J. and E.H.) and UG3NS123723 (S.D.), and Weill Neurohub (N.J.).

## METHODS

### Electrode fabrication

Electrode arrays were microfabricated using similar methods to previously described processes^101,102^. In brief, a photolithographic fabrication technique embedded Cr/Au bi-layer metal traces in a thin optically transparent parylene C substrate with PtAg discs at the end of each metal trace. Exposed PtAg electrode contact surfaces were dealloyed in hot nitric acid which left a platinum nanorod (PtNR) film. Two electrode array layouts using differently sized contacts (90 µm and 200 µm diameter) were fabricated and bonded to printed circuit boards (PCBs) using silver epoxy (MG Chemicals 8331)-based bump bonding, as described in a previous study^101^. Zero-insertion force (ZIF) connectors attached a ribbon cable between the electrode-bonded PCB and the stimulation/amplifier chip (Intan RHS headstage). Electrode impedances were measured at 1 kHz in saline using the Intan RHS Stim/Recording System and RHX software prior to mouse implantation to verify successful fabrication and electrical connection.

### Mouse surgery and electrode implantation

All animal experiments were conducted according to the National Institutes of Health guidelines for animal research. Procedures and protocols involving mice were approved by the Animal Care and Use Committee at the University of California, Berkeley.

Methods for chronic mouse implant surgery were adapted from previous studies^33,103^, with additional steps involving the PtNR electrode array. Mice aged 3-6 months, either wildtype (WT, JAX 000664) or transgenic with Cre-recombinase labeling of inhibitory cell subtypes (Pvalb-IRES-Cre, SST-IRES-Cre, NDNF-IRES-Cre; JAX 017320, 013044, and 030757, respectively), were anesthetized and head-fixed in a stereotaxic apparatus (Kopf Instruments). A 3.5-mm-diameter craniotomy was performed over the left primary visual cortex (V1, ∼2.5 mm medial-lateral and 1 mm anterior-posterior to lambda) and nine viral injections were performed in a 3 × 3 grid in the exposed cortex, with an average injection spacing of ∼600 µm. For WT animals, 30 nL of AAV2/1-syn-GCaMP6s at a titer of 1.3×10^13^ was injected at 250 µm below the brain surface per injection site. For PV-Cre, SST-Cre, and NDNF-Cre animals, 30 nL of AAV2/1-syn-FLEX-GCaMP6s at titers of 2.6×10^12^ to 1.3×10^13^ were injected at 250 µm below brain surface per injection site.

Cranial windows were made using two glass pieces of 170 µm thickness, a 3.5 mm diameter disk and a ring with 3 mm inner diameter, concentrically attached to each other with a UV-cured optical adhesive (Norland NOA61). The PtNR electrode array was then attached to the glass disk with the same optical adhesive, as demonstrated in a similar surgery^104^. The electrode array and cranial window were placed into the craniotomy and sealed with tissue adhesive (3M VetBond). Exposed electrode traces emerging from the posterior edge of the cranial window were coated with a surgical-grade silicone elastomer (Kwik-Sil, World Precision Instruments) to protect them from being damaged by dental acrylic, as demonstrated in previous chronic neural electrode implant methods^105,106^. A stainless-steel head-bar was then rigidly attached to the skull with dental acrylic and allowed to cure. The electrode ground was attached to the head-bar. A 3D resin-printed structure was attached to the head-bar and functioned as both the head-cone to reject stray light during 2PFM and as a housing for the electrode-bonded PCB (see Supplementary Materials for 3D design files). Finally, additional silicone elastomer, dental acrylic, and superglue (Loctite) were used to protect exposed electrode traces, provide additional structural support, and seal the implant, where needed. Implanted mice were provided with a post-operative analgesia (Meloxicam, SC, 5 mg/kg) for 2 days and allowed to recover for at least 2 weeks prior to experiments.

### In vivo imaging by 2PFM

Implanted mice were habituated to head fixation for 15 minutes approximately one week post-surgery, and again one or two days prior to their first imaging session. During imaging, mice were head-fixed under the microscope objective of a commercial 2PFM system with a Bessel imaging module^103^. All two-photon imaging was performed with 920 nm wavelength excitation light from a femtosecond titanium-sapphire laser (Chameleon Ultra II, Coherent Inc.). A Gaussian focus formed through a 25× 1.05-NA Olympus objective lens was used for fluorescence excitation for all data acquisition (post-objective power ranging between 4 and 89 mW: L1: 11.3±6.9 mW; L2/3: 26.5±12.0 mW; L4: 62.9±17.7 mW; mean ± S.D.), except for SST data, which was collected by scanning a 0.4-NA Bessel focus to improve throughput^46,107^ (post-objective power: 165 – 202 mW).

Blue drifting grating stimuli were displayed on a laptop screen (HP Spectre x360 13 with AMOLED screen) that was centered at approximately 10.5 cm from the mouse’s right eye and covered 75° × 75° of visual field. The gratings had 100% contrast, a spatial frequency of 0.07 cycles/degree, and temporal frequency of 2 cycles/s. A uniform visual stimulus with similar total luminance to that of the grating stimuli was presented to the mouse during baseline periods.

Retinotopic measurements were performed for each animal using a 10× 0.45 NA Nikon objective lens covering a FOV of 2.2 mm × 2.2 mm. The visual stimulation screen pseudorandomly flashed a bright square section of the screen chosen from a 3 × 3 grid spanning a total of 75° × 75° of visual field. By trial-averaging the calcium responses to each grid location, a retinotopic map was constructed to identify the region within the cranial window that corresponded to V1.

A 25× 1.05NA Olympus objective lens was used for high-resolution 2PFM imaging within V1 during concurrent electrical and/or visual stimulation. Image acquisition was performed at 15 Hz over a 781 × 781 µm FOV with 0.762 µm pixel size for 6 s per trial, beginning with 2 s of baseline followed by 4 s of stimulation response.

Electrically connected and low-impedance electrode contacts near the image FOV were chosen to be the source of electrical stimulation. Electrical stimuli, either alone or sharing the same onset time with grating stimuli, were pseudorandomly ordered and sent to the headstage via TCP control before every trial using a MATLAB (MathWorks) script. Electrode stimulation was triggered by a digital input from a photodiode reading the visual stimulation screen, to eliminate jitter caused by the screen’s refresh rate.

The onset of image acquisition was triggered using another photodiode reading the visual stimulation screen, such that the visual stimulation computer acted as the master clock for both 2PFM image acquisition and electrical stimulation. After the 6 s of image acquisition, there was a pause of 4 s for the fluorescence response to fall back to baseline levels, for a total of 10 seconds per trial. A total of 450 trials lasting 75 minutes consisted of 10 repeats of 45 unique trial types of visual-electrical stimulus pairs: 9 unique visual stimuli (8 drifting grating directions plus a blank control) and 5 unique electrical stimuli (cathode-leading or anode-leading 100 µA or 200 µA current strength, plus a 0 µA control). Animals were imaged up to 4 days total over a two-week period post-recovery, with each day consisting of up to 2 unique FOVs. In total, 62 unique FOVs were imaged across 14 animals split across 4 genetic backgrounds (WT: 3 animals, 14 FOVs, PV: 3 animals, 12 FOVs, SST: 5 animals, 21 FOVs, NDNF: 3 animals, 15 FOVs).

### Image preprocessing

All image preprocessing, visualization, and analyses were performed in ImageJ and MATLAB. Raw 2PFM images were first registered to remove rigid lateral motion artifacts. All ROIs were hand-drawn by the same person over cell bodies with the oval tool in ImageJ. A fluorescence trace, F_raw_(t), was extracted for each ROI by averaging the signals from all pixels within the ROI for each image frame.

We then subtracted neuropil contamination from the fluorescence traces. This step was especially critical for experiments such as ours since electrical stimulation directly and strongly excited neuropil. Specifically, fluorescence signal traces of neuropil surrounding each ROI were extracted and subtracted from F_raw_(t) following the procedure of a previous study^45^. The average fluorescence signal from a 35 µm radius area centered on each ROI, excluding pixels belonging to this or other ROIs, was extracted as the neuropil signal trace F_neuropil_(t). For each stimulus trial, the baseline fluorescence value F_0, neuropil_ was calculated from the 2-s baseline frames prior to stimulation onset, and the neuropil transient ΔF_neuropil_(t) = F_neuropil_(t) - F_0, neuropil_ was then calculated.

Neuropil subtraction coefficient ⍺ (**Supplementary Table 4**) was determined separately for each FOV, to account for differences in cell type-dependent neuropil labeling density, imaging method differences (Gaussian vs. Bessel beam), and imaging depth. For each ROI with a skewness value >2, we compared its F_neuropil_(t) with F_raw_(t) to find a neuropil subtraction coefficient ⍺ that best removed the contaminant neuropil signal (see Supplementary Fig. 2a from Ref. 45). We then calculated the average measured coefficient ⍺ in the FOV. We then calculated the neuropil-subtracted ROI fluorescence trace F_ROI_(t)=F_raw_(t) - ⍺ΔF_neuropil_(t). We set the baseline fluorescence F_0,ROI_ as the mode of the signal distribution, and calculated the calcium transient trace ΔF/F_0_(t)= (F_ROI_(t) - F_0,ROI_)/F_0,ROI_.

We removed all inactive ROIs from further analyses. ROIs were considered active if at any point during the experiment, fluorescence ΔF/F_0_ rose above the mean baseline + 3 standard deviations for 1 second. Baseline data were defined as the time points during the 2 seconds of data acquisition before the response period. Due to the large number of trials per experiment which resulted in extended 75-minute imaging sessions, we sometimes observed a slow drift in a ROI’s baseline over the course of the experiment. These slow drifts (on the order of minutes) could be due to several factors, including sample motion, brain state changes, and learning/habituation processes. To remove these effects from our data, for each ROI, we calculated the mean ΔF/F_0_ value from the 2-s baseline frames prior to stimulation onset, then subtracted it from ΔF/F_0_ of each trial. However, some ROIs showed highly varied baseline values within an experiment, so subtracting baselines from these ROIs would result in largely varied responses. Many of these ROIs with highly varied baselines had high rates of spontaneous activity and were not good ROI candidates to observe responses to electrical or visual stimulation. As a result, we removed ROIs with baseline brightness values that were very different from those of the ROI population (all ROIs of the same cell type): Specifically, baselines that were farther than the population median ± 3 times the median absolute deviation. This was a small proportion of ROIs (< 15%); their exclusion did not affect the findings of this study.

### Analysis and statistics

Electrical stimulation-only analyses were performed by analyzing only the subset of trials with blank visual stimulation (50 out of 450 trials). The ΔF/F_0_ traces of each ROI for all 10 trials of each electrical stimulation condition (0, 100_C_, 100_A_, 200_C_, 200_A_) were averaged to represent the ROI’s response to the stimulus. The population response was the averaged response of all individual ROIs. Shaded SEMs throughout this study were calculated based on the number of ROIs that were averaged. The “early response” ΔF/F_0_ for a given ROI was the average ΔF/F_0_ of the 15 frames (acquired at 15 Hz) beginning immediately post-stimulation, whereas the “late response” ΔF/F_0_ was the average of the last 15 frames of the trial (frames 46-60 post-stimulation). Statistical differences between the early responses of two electrical stimulation conditions were tested using two-sample Kolmogorov-Smirnov (KS) tests. 3D distances between ROIs and stimulating electrodes were calculated using the centers of both objects. Correlation strength and its statistical significance between an ROI’s early/late response strength and its 3D distance from the electrode were calculated by fitting a linear regression model.

ROIs with visually evoked activity were defined as ROIs that responded significantly to at least one visual stimulus. Specifically, an ROI’s late response to a visual stimulus in the absence of electrical stimulation was compared to its baseline period using a Student’s t-test. An ROI was considered to have visually evoked activity if at least one drifting grating direction tested significant against baseline (p < 0.05).

ROIs’ early responses to visual stimuli (in the absence of electrical stimulation) were fit to double gaussian curves to determine their preferred grating stimulus^108^:

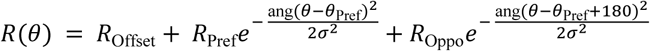

where *R*_Offset_ is a constant offset, and *R*_Pref_ and *R*_Oppo_ are the responses at grating angles *θ*_Pref_ and *θ*_Pref_ - 180°, respectively. The function ang(x) = min(|x|, |x−360|, |x+360|) wraps angular values onto the interval 0° to 180°. Global orientation selectivity indices (gOSI) were computed using the early response for grating angle *θ* as

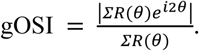

Statistical tests used are noted throughout the manuscript. p values are reported in the text or in Supplementary Tables.

---

**Supplementary Table 1:** Early response p-values of cell populations in different stimulation conditions.

| Visual stim conditions | E-stim conditions | Two-sample KS test p-values, EARLY RESPONSE |  |  |  |
| --- | --- | --- | --- | --- | --- |
|  |  | WT L2/3 | PV L2/3 | SST L2/3 | NDNF L1 |
| no vis stim | 200<sub>C</sub> vs. 0 | 1.7e-85 | 1.8e-01 | 2.1e-02 | 3.5e-08 |
|  | 100<sub>C</sub> vs. 0 | 8.3e-13 | 1.1e-02 | 5.6e-05 | 2.2e-07 |
|  | 100<sub>A</sub> vs. 0 | 2.3e-03 | 5.6e-06 | 2.1e-02 | 3.7e-06 |
|  | 200<sub>A</sub> vs. 0 | 7.5e-81 | 1.4e-04 | 1.2e-02 | 5.1e-09 |
|  | 200<sub>C</sub> vs. 200<sub>A</sub> | 1.0e-01 | 4.5e-02 | 4.4e-01 | 2.2e-01 |
|  | 100<sub>C</sub> vs. 100<sub>A</sub> | 2.5e-05 | 1.1e-01 | 6.9e-03 | 8.9e-01 |
| orthogonal | 200<sub>C</sub> vs. 0 | 3.2e-100 | 2.3e-02 | 2.6e-04 | 6.8e-10 |
|  | 100<sub>C</sub> vs. 0 | 8.6e-19 | 4.5e-01 | 4.4e-02 | 7.8e-05 |
|  | 100<sub>A</sub> vs. 0 | 1.9e-14 | 3.0e-01 | 2.3e-01 | 3.7e-06 |
|  | 200<sub>A</sub> vs. 0 | 1.4e-79 | 2.3e-02 | 5.6e-05 | 4.1e-12 |
|  | 200<sub>C</sub> vs. 200<sub>A</sub> | 3.3e-04 | 9.5e-01 | 8.6e-01 | 4.8e-01 |
|  | 100<sub>C</sub> vs. 100<sub>A</sub> | 1.1e-02 | 6.1e-02 | 6.5e-01 | 7.4e-01 |
| preferred | 200<sub>C</sub> vs. 0 | 2.6e-66 | 1.1e-24 | 1.4e-11 | 8.6e-13 |
|  | 100<sub>C</sub> vs. 0 | 9.9e-117 | 1.1e-13 | 3.9e-14 | 1.2e-17 |
|  | 100<sub>A</sub> vs. 0 | 1.3e-129 | 1.9e-11 | 2.8e-17 | 3.3e-14 |
|  | 200<sub>A</sub> vs. 0 | 6.2e-84 | 2.0e-17 | 1.4e-08 | 7.6e-14 |
|  | 200<sub>C</sub> vs. 200<sub>A</sub> | 4.3e-02 | 2.3e-01 | 5.8e-01 | 7.4e-01 |
|  | 100<sub>C</sub> vs. 100<sub>A</sub> | 3.3e-01 | 7.3e-01 | 8.6e-01 | 7.4e-01 |
| opposite | 200<sub>C</sub> vs. 0 | 4.1e-30 | 5.2e-03 | 5.3e-04 | 7.8e-05 |
|  | 100<sub>C</sub> vs. 0 | 2.3e-01 | 1.4e-01 | 7.5e-04 | 3.6e-02 |
|  | 100<sub>A</sub> vs. 0 | 3.7e-01 | 3.0e-01 | 1.2e-02 | 6.5e-02 |
|  | 200<sub>A</sub> vs. 0 | 9.4e-23 | 2.3e-03 | 8.3e-05 | 2.7e-02 |
|  | 200<sub>C</sub> vs. 200<sub>A</sub> | 2.5e-02 | 9.8e-01 | 8.6e-01 | 2.7e-01 |
|  | 100<sub>C</sub> vs. 100<sub>A</sub> | 9.9e-01 | 4.5e-02 | 4.4e-01 | 9.9e-01 |

**Supplementary Table 2:**
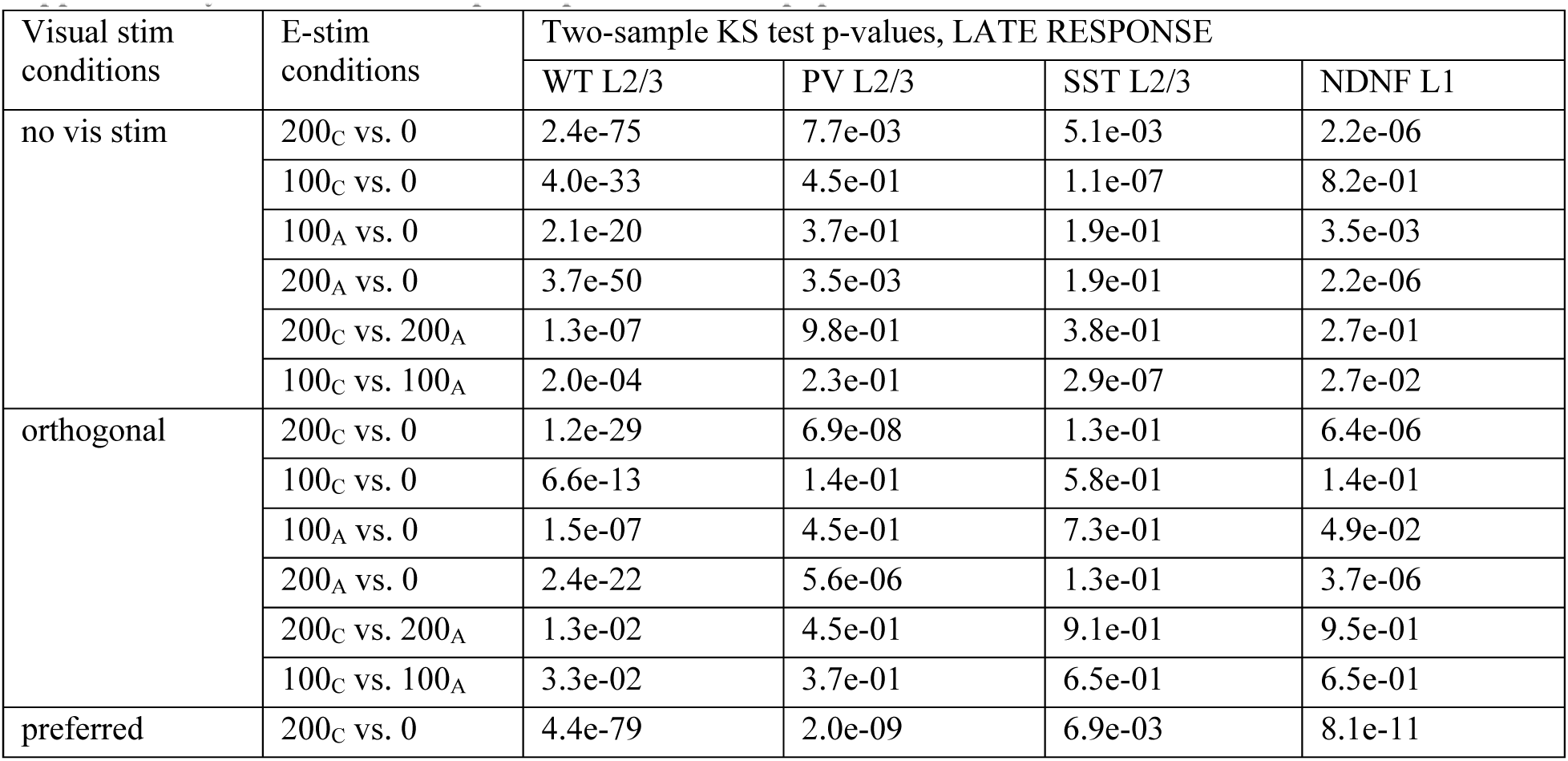
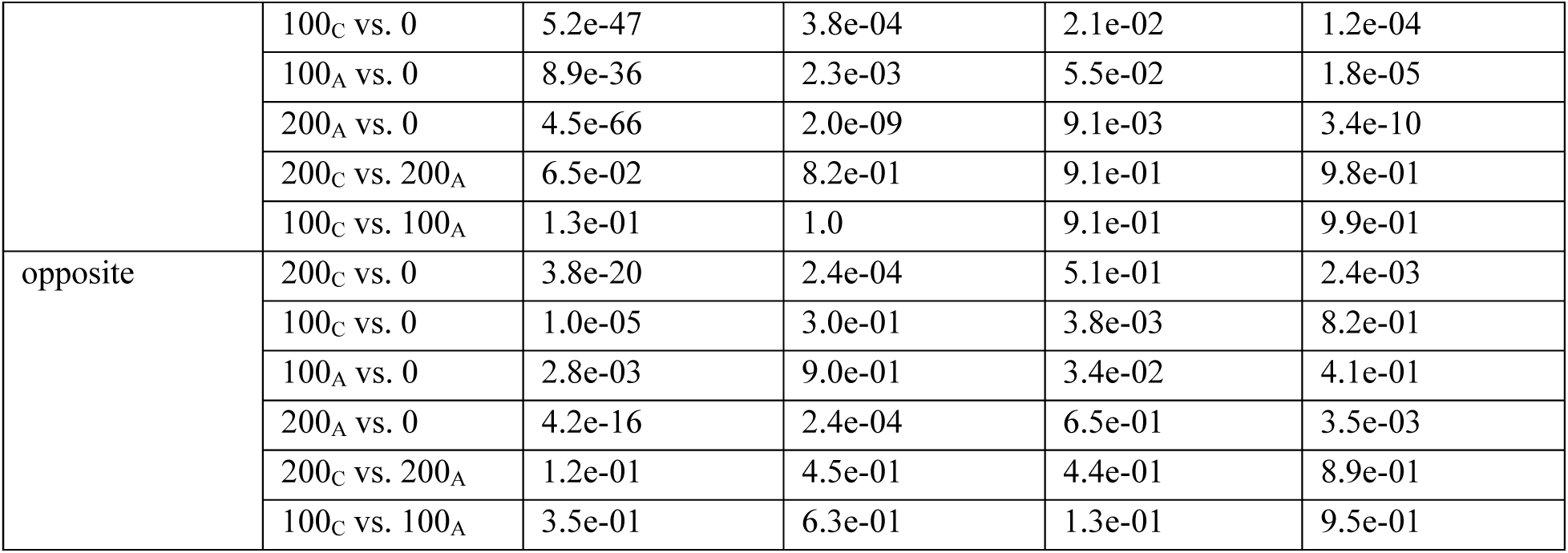
Late response p-values of cell populations in different stimulation conditions.

| Visual stim conditions | E-stim conditions | Two-sample KS test p-values, LATE RESPONSE |  |  |  |
| --- | --- | --- | --- | --- | --- |
|  |  | WT L2/3 | PV L2/3 | SST L2/3 | NDNF L1 |
| no vis stim | 200<sub>C</sub> vs. 0 | 2.4e-75 | 7.7e-03 | 5.1e-03 | 2.2e-06 |
|  | 100<sub>C</sub> vs. 0 | 4.0e-33 | 4.5e-01 | 1.1e-07 | 8.2e-01 |
|  | 100<sub>A</sub> vs. 0 | 2.1e-20 | 3.7e-01 | 1.9e-01 | 3.5e-03 |
|  | 200<sub>A</sub> vs. 0 | 3.7e-50 | 3.5e-03 | 1.9e-01 | 2.2e-06 |
|  | 200<sub>C</sub> vs. 200<sub>A</sub> | 1.3e-07 | 9.8e-01 | 3.8e-01 | 2.7e-01 |
|  | 100<sub>C</sub> vs. 100<sub>A</sub> | 2.0e-04 | 2.3e-01 | 2.9e-07 | 2.7e-02 |
| orthogonal | 200<sub>C</sub> vs. 0 | 1.2e-29 | 6.9e-08 | 1.3e-01 | 6.4e-06 |
|  | 100<sub>C</sub> vs. 0 | 6.6e-13 | 1.4e-01 | 5.8e-01 | 1.4e-01 |
|  | 100<sub>A</sub> vs. 0 | 1.5e-07 | 4.5e-01 | 7.3e-01 | 4.9e-02 |
|  | 200<sub>A</sub> vs. 0 | 2.4e-22 | 5.6e-06 | 1.3e-01 | 3.7e-06 |
|  | 200<sub>C</sub> vs. 200<sub>A</sub> | 1.3e-02 | 4.5e-01 | 9.1e-01 | 9.5e-01 |
|  | 100<sub>C</sub> vs. 100<sub>A</sub> | 3.3e-02 | 3.7e-01 | 6.5e-01 | 6.5e-01 |
| preferred | 200<sub>C</sub> vs. 0 | 4.4e-79 | 2.0e-09 | 6.9e-03 | 8.1e-11 |
|  | 100<sub>C</sub> vs. 0 | 5.2e-47 | 3.8e-04 | 2.1e-02 | 1.2e-04 |
|  | 100<sub>A</sub> vs. 0 | 8.9e-36 | 2.3e-03 | 5.5e-02 | 1.8e-05 |
|  | 200<sub>A</sub> vs. 0 | 4.5e-66 | 2.0e-09 | 9.1e-03 | 3.4e-10 |
|  | 200<sub>C</sub> vs. 200<sub>A</sub> | 6.5e-02 | 8.2e-01 | 9.1e-01 | 9.8e-01 |
|  | 100<sub>C</sub> vs. 100<sub>A</sub> | 1.3e-01 | 1.0 | 9.1e-01 | 9.9e-01 |
| opposite | 200<sub>C</sub> vs. 0 | 3.8e-20 | 2.4e-04 | 5.1e-01 | 2.4e-03 |
|  | 100<sub>C</sub> vs. 0 | 1.0e-05 | 3.0e-01 | 3.8e-03 | 8.2e-01 |
|  | 100<sub>A</sub> vs. 0 | 2.8e-03 | 9.0e-01 | 3.4e-02 | 4.1e-01 |
|  | 200<sub>A</sub> vs. 0 | 4.2e-16 | 2.4e-04 | 6.5e-01 | 3.5e-03 |
|  | 200<sub>C</sub> vs. 200<sub>A</sub> | 1.2e-01 | 4.5e-01 | 4.4e-01 | 8.9e-01 |
|  | 100<sub>C</sub> vs. 100<sub>A</sub> | 3.5e-01 | 6.3e-01 | 1.3e-01 | 9.5e-01 |

**Supplementary Table 3:** p values for statistical testing of gOSI distributions.

| Wilcoxon ranksum test p-values | gOSI, Fig. 2h |
| --- | --- |
| 200<sub>C</sub> < 0 | 4.9e-54 |
| 100<sub>C</sub> < 0 | 9.7e-10 |
| 100<sub>A</sub> < 0 | 5.6e-5 |
| 200<sub>A</sub> < 0 | 8.3e-30 |
| 200<sub>C</sub> < 100<sub>C</sub> | 5.2e-34 |
| 200<sub>A</sub> < 100<sub>A</sub> | 6.0e-24 |

**Supplementary Table 4:** Neuropil subtraction coefficient ⍺.

| Neuropil subtraction coefficient | Number of FOVs | Mean | Std |
| --- | --- | --- | --- |
| WT | 14 | 0.74 | 0.05 |
| PV L2/3 | 6 | 0.36 | 0.11 |
| PV L4 | 6 | 0.41 | 0.14 |
| SST L2/3 (Bessel) | 8 | 0.81 | 0.07 |
| SST L4 | 12 | 0.24 | 0.14 |
| NDNF | 15 | 0.24 | 0.22 |

**Supplementary Figure 1.**
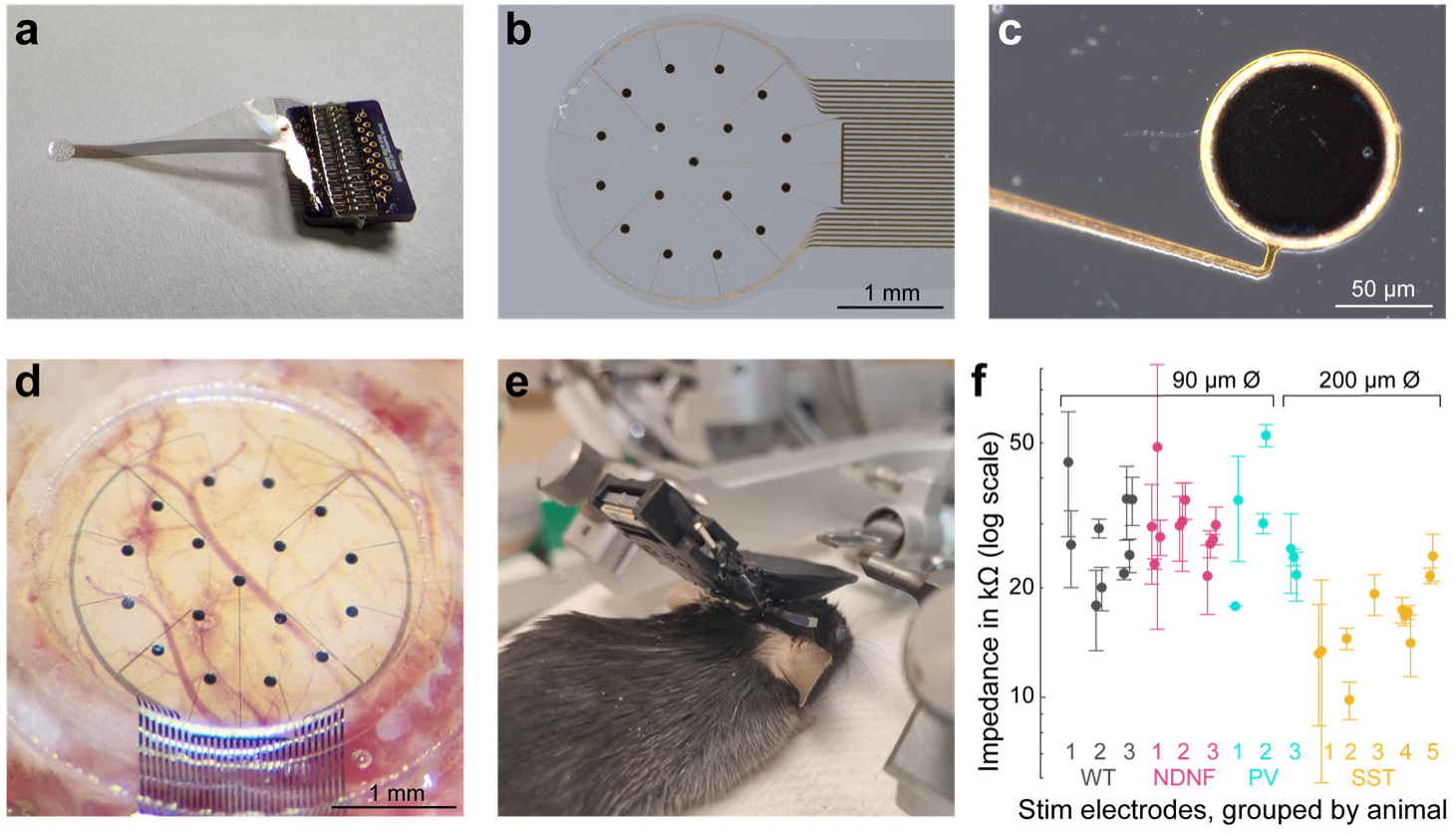
Chronic implantation of a PtNR ECoG electrode array in mice. (**a**) PCB-bonded flexible microelectrode array. (**b**) Zoomed-in view of electrode array. (**c**) Further zoomed-in view of a single electrode contact. (**d**) Electrode glued to a glass window, implanted in a craniotomy during surgery. (**e**) A mouse post surgery, with a 3D-printed implant housing and head bar. (**f**) Impedances of electrode contacts used for electrical stimulation for each mouse. Each circle represents one unique electrode contact measured post implantation. Two electrode contact diameters were used in this study, as labeled. Error bars: S.D. of impedance for each electrode contact.

**Supplementary Figure 2.**
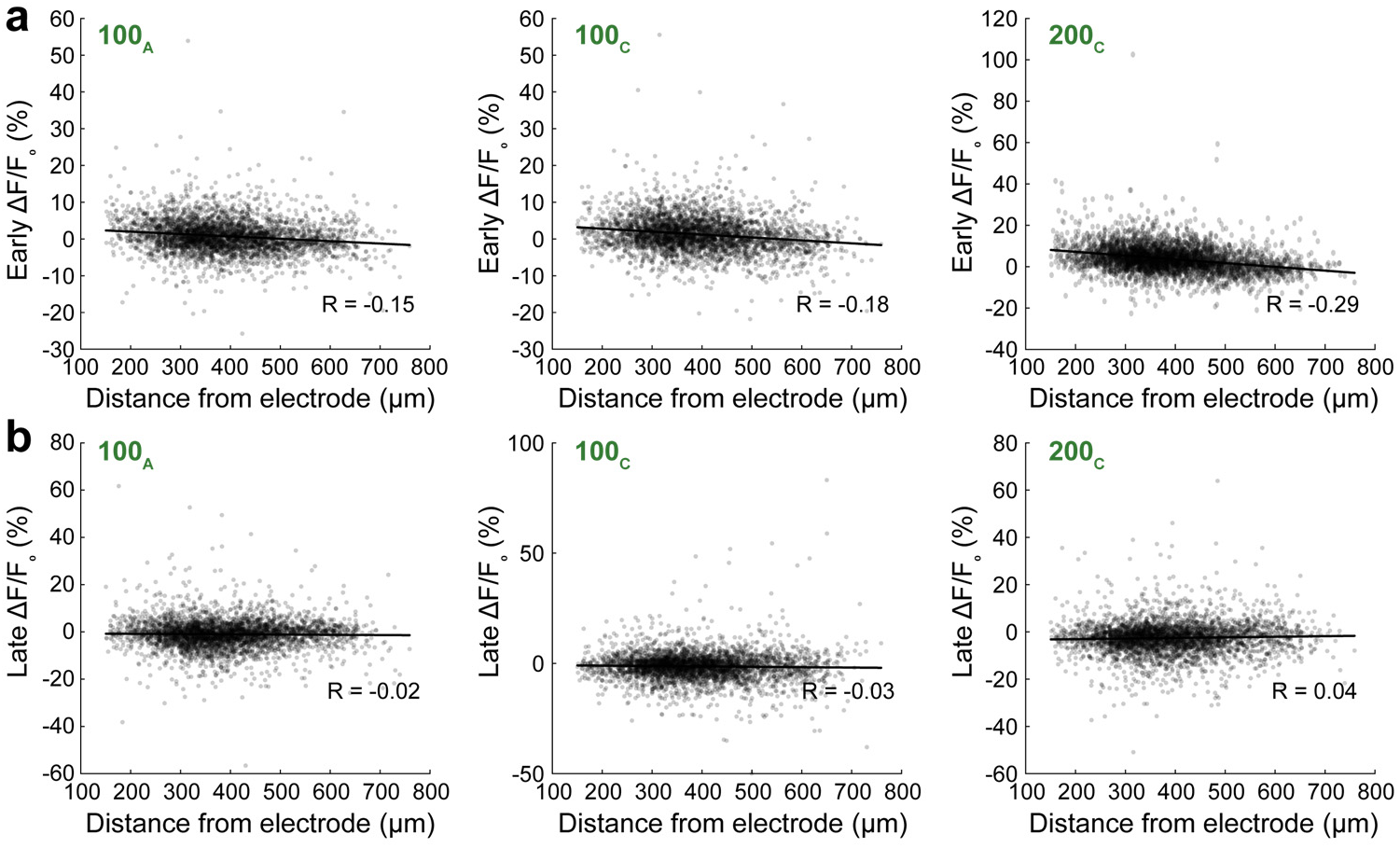
Early and late responses of L2/3 neurons versus their 3D distance from electrode center. (**a**) Scatter plots of early and (**b**) late responses of each L2/3 neurons to 100_A_, 100_C_, and 200_C_ E-Stim vs. its 3D distance from electrode center. Two-tailed test for Pearson’s correlation coefficient, p values: (**a**) 1.1×10^−20^, 5.0×10^−27^, 1.8×10^−75^ for early response to 100_A_, 100_C_, and 200_C_ E-Stim, respectively; (**b**) 0.29, 0.09, 0.02 for late response to 100_A_, 100_C_, and 200_C_ E-Stim, respectively.

**Supplementary Figure 3.**
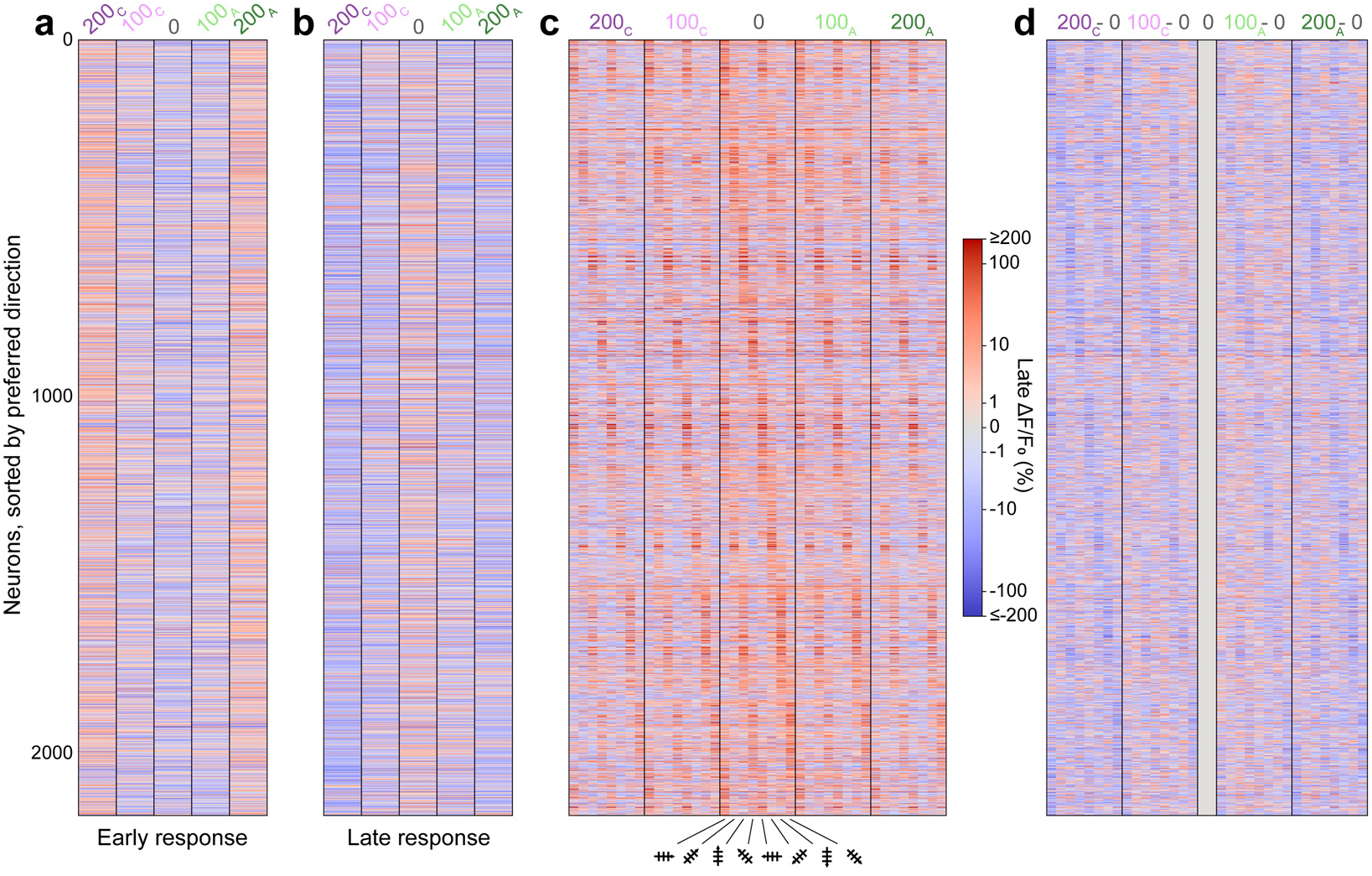
Early and late ΔF/F0 for individual L2/3 neurons. Same data as in Fig. 2. (**a,b**) Heatmaps showing the early response ΔF/F_0_ (**a**) and late response ΔF/F_0_ (**b**) of 2,173 L2/3 neurons under E-Stim. Rows: individual cells sorted in the same way as in Fig. 2d; Columns: electrical stimulation conditions. (**c**) Heatmap showing the late response ΔF/F_0_ of 2,173 L2/3 neurons under V-Stim and EV-Stim. Rows: individual cells sorted by their preferred direction; Columns: electrical-visual stimulation conditions. (**d**) Heatmap showing the difference in late response ΔF/F_0_, by subtracting V-Stim responses from EV-Stim responses in **c**. Blue: reduced activity; red: increased activity.

**Supplementary Figure 4.**
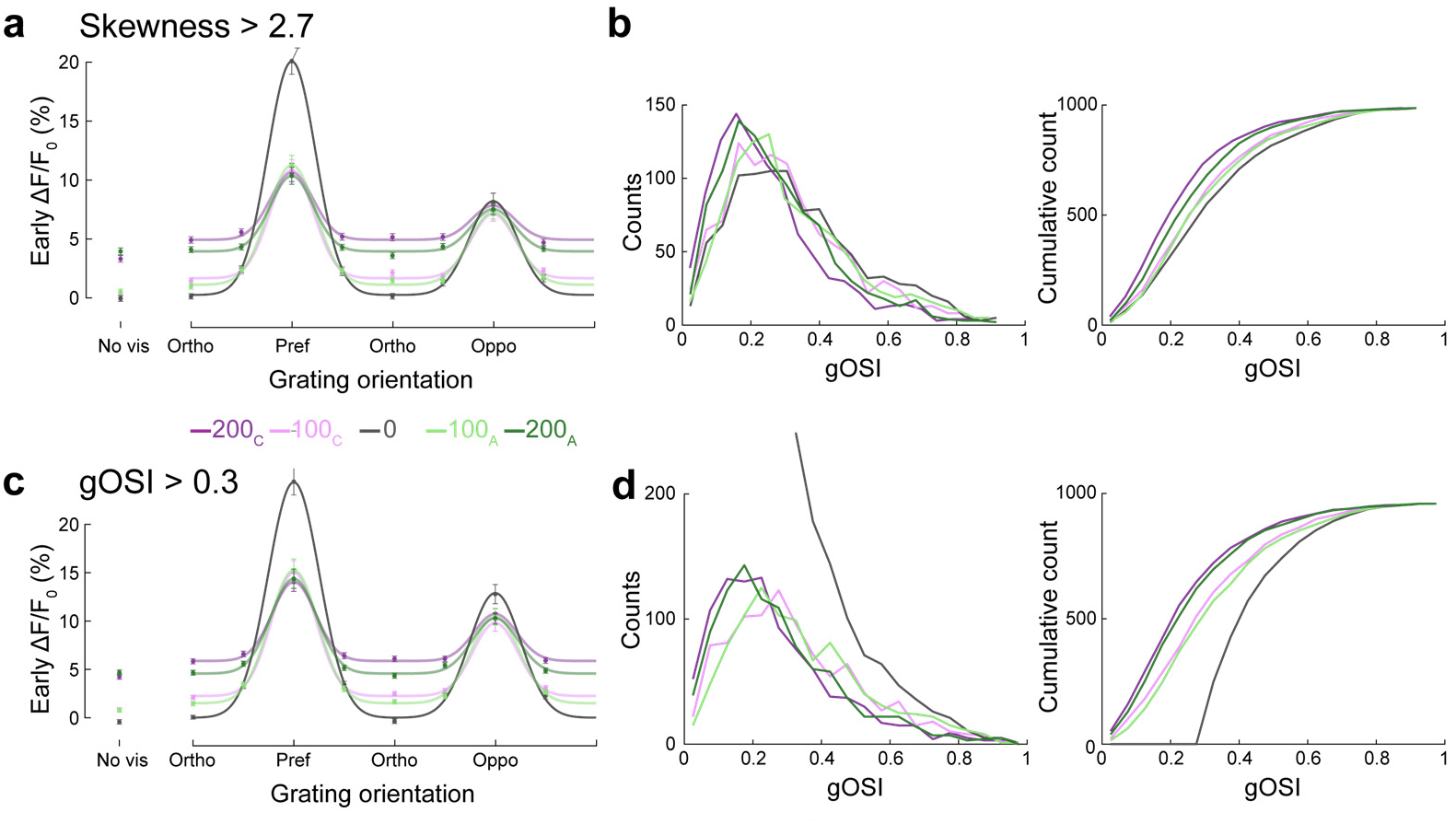
Electrical stimulation reduces orientation selectivity in putative L2/3 pyramidal neurons. **(a,b)** Similar analyses to Fig. 2f**,h** on 986 L2/3 neurons with visually evoked activity and fluorescence trace skewness values > 2.7. **(c,d**) Analyses on 959 L2/3 neurons with visually evoked activity and gOSI > 0.3. (**a,c**) Population-averaged early responses for drifting grating stimuli and the fitted orientation tuning curves under 5 electrical stimulation conditions. (**b,d**) Distributions and cumulative distributions of gOSI values under 5 electrical stimulation conditions.

**Supplementary Figure 5.**
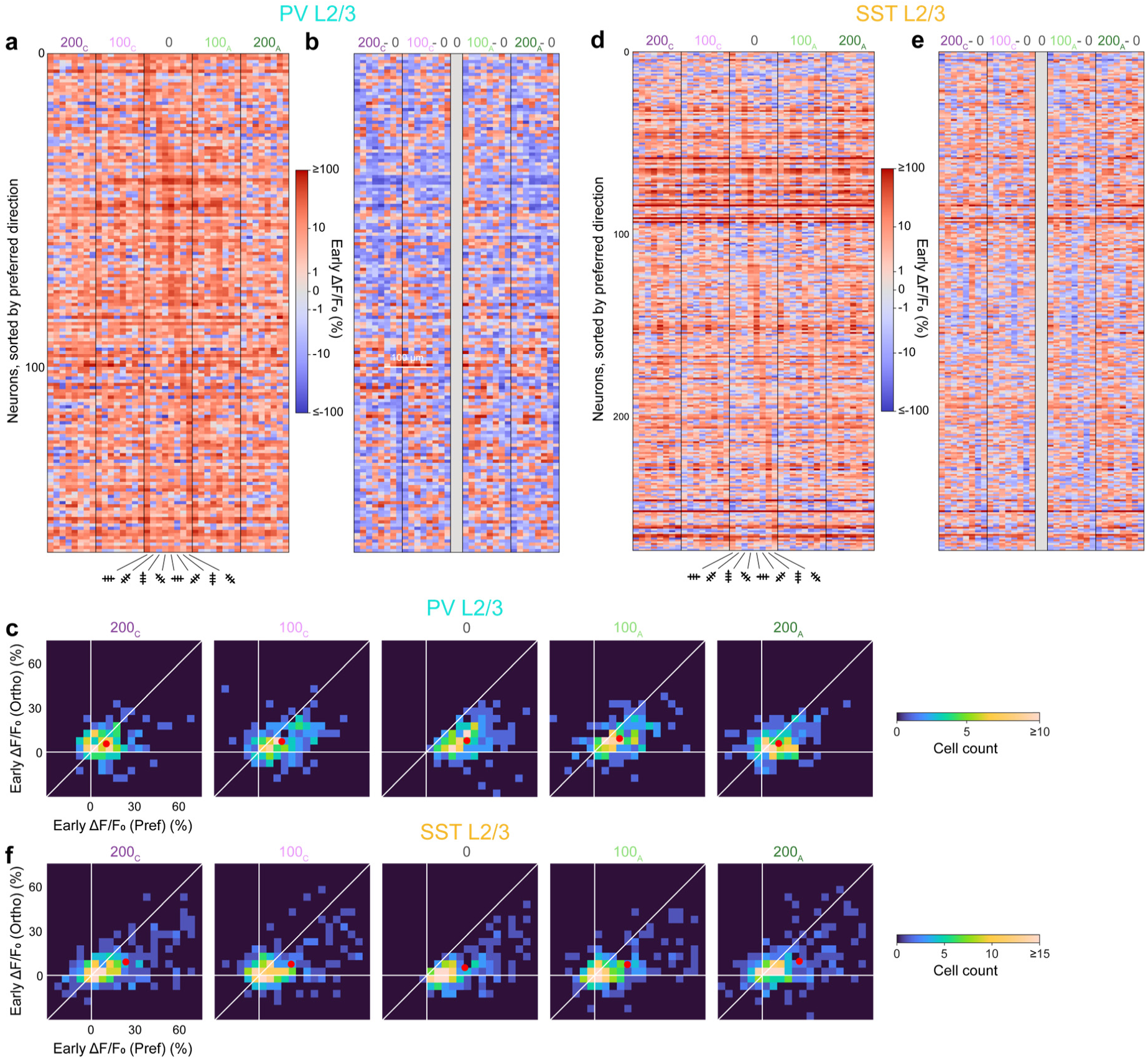
Early response of individual PV and SST neurons INs. Same data as in Fig. 3. (**a**) Heatmap showing the early response ΔF/F_0_ of 156 L2/3 PV INs. Rows: individual cells sorted by their preferred direction; Columns: electrical-visual stimulation conditions. (**b**) Heatmap showing the difference in early response ΔF/F_0_, by subtracting V-Stim responses from EV-Stim responses in **a**. Blue: reduction in response; red: increase in response. (**c**) Neurons’ early response towards pref vs. ortho gratings, under 5 electrical stimulation conditions. Red dots: mean responses. (**d,e,f**) Same as **a,b,c**, but for 274 SST INs.

**Supplementary Figure 6.**
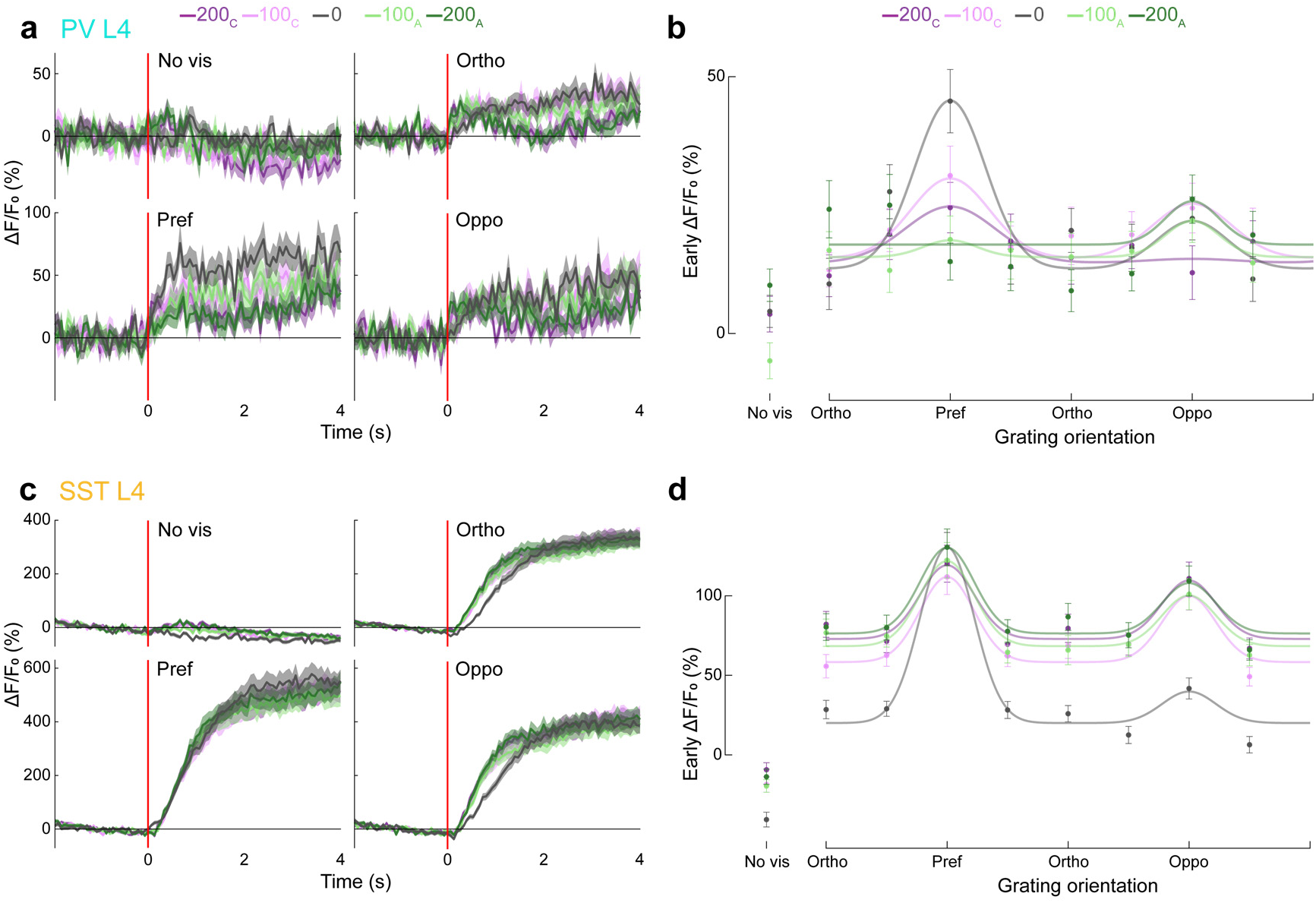
L4 PV and SST INs respond similarly to L2/3 PV and SST INs. **b**) Population- and trial-averaged ΔF/F_0_ traces of 133 L4 PV INs with visually-evoked activity under 5 electrical stimulation conditions, grouped for 4 visual stimulation conditions: no stimulus, ortho, pref, and oppo. Trace and shade: mean and SEM. Red lines: stimulus onset. (**b**) Scattered data: early-response ΔF/F_0_ of the traces in **a**; tuning curves: double-Gaussian fits to scattered data. (**c,d**) Same as **a,b**, but for 413 L4 SST INs with visually-evoked activity.

**Supplementary Figure 7.**
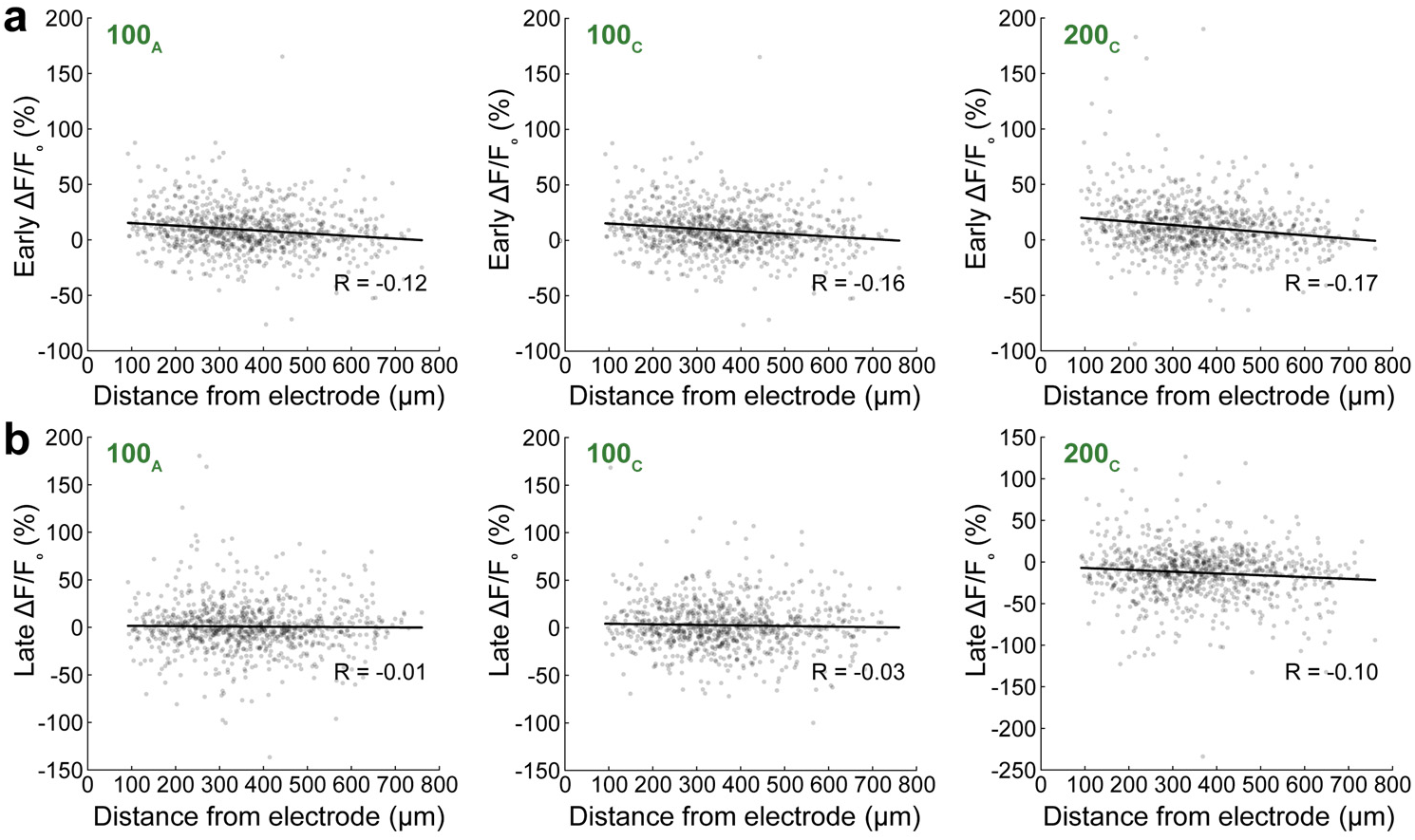
Early and late responses of NDNF neurons versus 3D distance from electrode center. (**a**) Scatter plots of early and (**b**) late responses of each NDNF neurons to 100_A_, 100_C_, and 200_C_ E-Stim vs. its 3D distance from electrode center. Two-tailed test for Pearson’s correlation coefficient, p values: (**a**) 3.4×10^−4^, 2.3×10^−6^, 1.3×10^−7^ for early response to 100_A_, 100_C_, and 200_C_ E-Stim, respectively; (**b**) 0.7, 0.34, 0.0042 for late response to 100_A_, 100_C_, and 200_C_ E-Stim, respectively

## References

1 Moon, H. et al. Electrocorticogram (ECoG): Engineering approaches and clinical challenges for translational medicine. *Adv*. Mater. Technol. 9, 2301692, doi:10.1002/admt.202301692 (2024).

2 Yang, T., Hakimian, S. & Schwartz, T. H. Intraoperative ElectroCorticoGraphy (ECog): indications, techniques, and utility in epilepsy surgery. Epileptic Disord. 16, 271–279, doi:10.1684/epd.2014.0675 (2014).

3 Vakani, R. & Nair, D. R. Electrocorticography and functional mapping. Handb. Clin. Neurol. 160, 313–327, doi:10.1016/B978-0-444-64032-1.00020-5 (2019).

4 Penfield, W. & Jasper, H. H. Epilepsy and the functional anatomy of the human brain. [1st ed.] edn, (Little, Brown, 1954).

5 Penfield, W. & Perot, P. The brain’s record of auditory and visual experience. Brain 86, 595–696, doi:10.1093/brain/86.4.595 (1963).

6 Ojemann, G., Ojemann, J., Lettich, E. & Berger, M. Cortical language localization in left, dominant hemisphere. An electrical stimulation mapping investigation in 117 patients. J. Neurosurg. 71, 316–326, doi:10.3171/jns.1989.71.3.0316 (1989).

7 Degenhart, A. D. et al. Histological evaluation of a chronically-implanted electrocorticographic electrode grid in a non-human primate. J. Neural Eng. 13, 046019, doi:10.1088/1741-2560/13/4/046019 (2016).

8 Ryapolova-Webb, E. et al. Chronic cortical and electromyographic recordings from a fully implantable device: preclinical experience in a nonhuman primate. J. Neural Eng. 11, 016009, doi:10.1088/1741-2560/11/1/016009 (2014).

9 Fifer, M. S. et al. Intracortical somatosensory stimulation to elicit fingertip sensations in an individual with spinal cord injury. Neurology 98, e679–e687, doi:10.1212/WNL.0000000000013173 (2022).

10 Lee, B. et al. Engineering artificial somatosensation through cortical stimulation in humans. Front. Syst. Neurosci. 12, 24, doi:10.3389/fnsys.2018.00024 (2018).

11 Lewis, P. M. & Rosenfeld, J. V. Electrical stimulation of the brain and the development of cortical visual prostheses: An historical perspective. Brain Res. 1630, 208–224, doi:10.1016/j.brainres.2015.08.038 (2016).

12 Tehovnik, E. J., Slocum, W. M. & Schiller, P. H. Differential effects of laminar stimulation of V1 cortex on target selection by macaque monkeys: Stimulation of V1 affects target selection. Eur. J. Neurosci. 16, 751–760, doi:10.1046/j.1460-9568.2002.02123.x (2002).

13 Salzman, C. D., Britten, K. H. & Newsome, W. T. Cortical microstimulation influences perceptual judgements of motion direction. Nature 346, 174–177, doi:10.1038/346174a0 (1990).

14 Halgren, E., Walter, R. D., Cherlow, D. G. & Crandall, P. H. Mental phenomena evoked by electrical stimulation of the human hippocampal formation and amygdala. Brain 101, 83–117, doi:10.1093/brain/101.1.83 (1978).

15 Blumenfeld, Z. & Brontë-Stewart, H. High frequency deep brain stimulation and neural rhythms in Parkinson’s disease. Neuropsychol. Rev. 25, 384–397, doi:10.1007/s11065-015-9308-7 (2015).

16 Papageorgiou, P. N., Deschner, J. & Papageorgiou, S. N. Effectiveness and adverse effects of deep brain stimulation: Umbrella review of meta-analyses. J. Neurol. Surg. A Cent. Eur. Neurosurg. 78, 180–190, doi:10.1055/s-0036-1592158 (2017).

17 Baizabal-Carvallo, J. F. & Alonso-Juarez, M. Low-frequency deep brain stimulation for movement disorders. Parkinsonism Relat. Disord. 31, 14–22, doi:10.1016/j.parkreldis.2016.07.018 (2016).

18 Nagaraj, V. et al. Future of seizure prediction and intervention: closing the loop: Closing the loop. J. Clin. Neurophysiol. 32, 194–206, doi:10.1097/WNP.0000000000000139 (2015).

19 Hummel, F. C. & Cohen, L. G. Non-invasive brain stimulation: a new strategy to improve neurorehabilitation after stroke? Lancet Neurol. 5, 708–712, doi:10.1016/S1474-4422(06)70525-7 (2006).

20 Stoney, S. D., Jr., Thompson, W. D. & Asanuma, H. Excitation of pyramidal tract cells by intracortical microstimulation: effective extent of stimulating current. J. Neurophysiol. 31, 659–669, doi:10.1152/jn.1968.31.5.659 (1968).

21 Histed, M. H., Bonin, V. & Reid, R. C. Direct activation of sparse, distributed populations of cortical neurons by electrical microstimulation. Neuron 63, 508–522, doi:10.1016/j.neuron.2009.07.016 (2009).

22 Ranck, J. B., Jr. Which elements are excited in electrical stimulation of mammalian central nervous system: a review. Brain Res. 98, 417–440, doi:10.1016/0006-8993(75)90364-9 (1975).

23 Nowak, L. G. & Bullier, J. Axons, but not cell bodies, are activated by electrical stimulation in cortical gray matter. I. Evidence from chronaxie measurements. Exp. Brain Res. 118, 477–488, doi:10.1007/s002210050304 (1998).

24 Rattay, F. The basic mechanism for the electrical stimulation of the nervous system. Neuroscience 89, 335–346, doi:10.1016/s0306-4522(98)00330-3 (1999).

25 Michelson, N. J., Eles, J. R., Vazquez, A. L., Ludwig, K. A. & Kozai, T. D. Y. Calcium activation of cortical neurons by continuous electrical stimulation: Frequency dependence, temporal fidelity, and activation density. J. Neurosci. Res. 97, 620–638, doi:10.1002/jnr.24370 (2019).

26 Tehovnik, E. J., Tolias, A. S., Sultan, F., Slocum, W. M. & Logothetis, N. K. Direct and Indirect Activation of Cortical Neurons by Electrical Microstimulation. Journal of Neurophysiology 96, 512–521, doi:10.1152/jn.00126.2006 (2006).

27 Eles, J. R., Stieger, K. C. & Kozai, T. D. Y. The temporal pattern of intracortical microstimulation pulses elicits distinct temporal and spatial recruitment of cortical neuropil and neurons. J. Neural Eng. 18, 015001, doi:10.1088/1741-2552/abc29c (2021).

28 Eles, J. R. & Kozai, T. D. Y. In vivo imaging of calcium and glutamate responses to intracortical microstimulation reveals distinct temporal responses of the neuropil and somatic compartments in layer II/III neurons. Biomaterials 234, 119767, doi:10.1016/j.biomaterials.2020.119767 (2020).

29 Dadarlat, M. C., Sun, Y. J. & Stryker, M. P. Activity-dependent recruitment of inhibition and excitation in the awake mammalian cortex during electrical stimulation. Neuron 112, 821–834.e824, doi:10.1016/j.neuron.2023.11.022 (2024).

30 Neumann, W.-J., Steiner, L. A. & Milosevic, L. Neurophysiological mechanisms of deep brain stimulation across spatiotemporal resolutions. Brain 146, 4456–4468, doi:10.1093/brain/awad239 (2023).

31 Chen, T.-W. et al. Ultrasensitive fluorescent proteins for imaging neuronal activity. Nature 499, 295–300, doi:10.1038/nature12354 (2013).

32 Fan, J. L. et al. High-speed volumetric two-photon fluorescence imaging of neurovascular dynamics. Nat. Commun. 11, 6020, doi:10.1038/s41467-020-19851-1 (2020).

33 Sun, W., Tan, Z., Mensh, B. D. & Ji, N. Thalamus provides layer 4 of primary visual cortex with orientation- and direction-tuned inputs. Nat. Neurosci. 19, 308–315, doi:10.1038/nn.4196 (2016).

34 Hippenmeyer, S. et al. A developmental switch in the response of DRG neurons to ETS transcription factor signaling. PLoS Biol. 3, e159, doi:10.1371/journal.pbio.0030159 (2005).

35 Madisen, L. et al. A robust and high-throughput Cre reporting and characterization system for the whole mouse brain. Nat. Neurosci. 13, 133–140, doi: 10.1038/nn.2467 (2010).

36 Taniguchi, H. et al. A Resource of Cre Driver Lines for Genetic Targeting of GABAergic Neurons in Cerebral Cortex. Neuron 71, 995–1013, doi:10.1016/j.neuron.2011.07.026 (2011).

37 Schuman, B. et al. Four Unique Interneuron Populations Reside in Neocortical Layer 1. J. Neurosci. 39, 125–139, doi:10.1523/JNEUROSCI.1613-18.2018 (2019).

38 Lu, R. et al. Rapid mesoscale volumetric imaging of neural activity with synaptic resolution. Nat. Methods 17, 291–294, doi:10.1038/s41592-020-0760-9 (2020).

39 Lu, R. et al. Video-rate volumetric functional imaging of the brain at synaptic resolution. bioRxiv, doi:10.1101/058495 (2016).

40 Carandini, M. & Heeger, D. J. Summation and division by neurons in primate visual cortex. Science 264, 1333–1336, doi:10.1126/science.8191289 (1994).

41 Wilson, N. R., Runyan, C. A., Wang, F. L. & Sur, M. Division and subtraction by distinct cortical inhibitory networks in vivo. Nature 488, 343–348, doi:10.1038/nature11347 (2012).

42 Kreile, A. K., Bonhoeffer, T. & Hübener, M. Altered visual experience induces instructive changes of orientation preference in mouse visual cortex. J. Neurosci. 31, 13911–13920, doi:10.1523/JNEUROSCI.2143-11.2011 (2011).

43 Xu, X., Roby, K. D. & Callaway, E. M. Immunochemical characterization of inhibitory mouse cortical neurons: three chemically distinct classes of inhibitory cells. J. Comp. Neurol. 518, 389–404, doi:10.1002/cne.22229 (2010).

44 Tremblay, R., Lee, S. & Rudy, B. GABAergic Interneurons in the Neocortex: From Cellular Properties to Circuits. Neuron 91, 260–292, doi:10.1016/j.neuron.2016.06.033 (2016).

45 Dipoppa, M. et al. Vision and Locomotion Shape the Interactions between Neuron Types in Mouse Visual Cortex. Neuron 98, 602–615.e608, doi:10.1016/j.neuron.2018.03.037 (2018).

46 Lu, R. et al. Video-rate volumetric functional imaging of the brain at synaptic resolution. Nat. Neurosci. 20, 620–628, doi:10.1038/nn.4516 (2017).

47 Abs, E. et al. Learning-related plasticity in dendrite-targeting layer 1 interneurons. Neuron 100, 684–699.e686, doi:10.1016/j.neuron.2018.09.001 (2018).

48 Anastasiades, P. G., Collins, D. P. & Carter, A. G. Mediodorsal and ventromedial thalamus engage distinct L1 circuits in the prefrontal cortex. Neuron 109, 314–330.e314, doi:10.1016/j.neuron.2020.10.031 (2021).

49 Cohen-Kashi Malina, K., et al. NDNF interneurons in layer 1 gain-modulate whole cortical columns according to an animal’s behavioral state. Neuron 109, 2150–2164.e2155, doi:10.1016/j.neuron.2021.05.001 (2021).

50 Naumann, L. B., Hertäg, L., Müller, J., Letzkus, J. J. & Sprekeler, H. Layer-specific control of inhibition by NDNF interneurons. Proc. Natl. Acad. Sci. U. S. A. 122, e2408966122, doi:10.1073/pnas.2408966122 (2025).

51 Schneider-Mizell, C. M. et al. Inhibitory specificity from a connectomic census of mouse visual cortex. Nature 640, 448–458, doi:10.1038/s41586-024-07780-8 (2025).

52 Oláh, S. et al. Regulation of cortical microcircuits by unitary GABA-mediated volume transmission. Nature 461, 1278–1281, doi:10.1038/nature08503 (2009).

53 Huang, S., Wu, S. J., Sansone, G., Ibrahim, L. A. & Fishell, G. Layer 1 neocortex: Gating and integrating multidimensional signals. Neuron 112, 184–200, doi:10.1016/j.neuron.2023.09.041 (2024).

54 Peng, H. et al. Morphological diversity of single neurons in molecularly defined cell types. Nature 598, 174–181, doi:10.1038/s41586-021-03941-1 (2021).

55 Stieger, K. C., Eles, J. R., Ludwig, K. A. & Kozai, T. D. Y. In vivo microstimulation with cathodic and anodic asymmetric waveforms modulates spatiotemporal calcium dynamics in cortical neuropil and pyramidal neurons of male mice. J. Neurosci. Res. 98, 2072–2095, doi:10.1002/jnr.24676 (2020).

56 Stieger, K. C., Eles, J. R., Ludwig, K. A. & Kozai, T. D. Y. Intracortical microstimulation pulse waveform and frequency recruits distinct spatiotemporal patterns of cortical neuron and neuropil activation. J. Neural Eng. 19, 026024, doi:10.1088/1741-2552/ac5bf5 (2022).

57 Komarov, M. et al. Selective recruitment of cortical neurons by electrical stimulation. PLoS Comput. Biol. 15, e1007277, doi:10.1371/journal.pcbi.1007277 (2019).

58 Kubota, Y. Untangling GABAergic wiring in the cortical microcircuit. Curr. Opin. Neurobiol. 26, 7–14, doi:10.1016/j.conb.2013.10.003 (2014).

59 Muñoz, W., Tremblay, R., Levenstein, D. & Rudy, B. Layer-specific modulation of neocortical dendritic inhibition during active wakefulness. Science 355, 954–959, doi:10.1126/science.aag2599 (2017).

60 Scala, F. et al. Layer 4 of mouse neocortex differs in cell types and circuit organization between sensory areas. Nat. Commun. 10, 4174, doi:10.1038/s41467-019-12058-z (2019).

61 Voigt, M. B. & Kral, A. Cathodic-leading pulses are more effective than anodic-leading pulses in intracortical microstimulation of the auditory cortex. J. Neural Eng. 16, 036002, doi:10.1088/1741-2552/ab0944 (2019).

62 Soh, D., Ten Brinke, T. R., Lozano, A. M. & Fasano, A. Therapeutic window of deep brain stimulation using cathodic monopolar, bipolar, semi-bipolar, and anodic stimulation. Neuromodulation 22, 451–455, doi:10.1111/ner.12957 (2019).

63 Kirsch, A. D., Hassin-Baer, S., Matthies, C., Volkmann, J. & Steigerwald, F. Anodic versus cathodic neurostimulation of the subthalamic nucleus: A randomized-controlled study of acute clinical effects. Parkinsonism Relat. Disord. 55, 61–67, doi:10.1016/j.parkreldis.2018.05.015 (2018).

64 Jiang, X. et al. Principles of connectivity among morphologically defined cell types in adult neocortex. Science 350, doi:10.1126/science.aac9462 (2015).

65 Swadlow, H. A. Antidromic activation: Measuring the refractory period at the site of axonal stimulation. Experimental Neurology 75, 514–519, doi:10.1016/0014-4886(82)90179-0 (1982).

66 Gamlin, C. R. et al. Connectomics of predicted Sst transcriptomic types in mouse visual cortex. Nature 640, 497–505, doi:10.1038/s41586-025-08805-6 (2025).

67 Johnson, L. A. et al. Direct electrical stimulation of the somatosensory cortex in humans using electrocorticography electrodes: a qualitative and quantitative report. J. Neural Eng. 10, 036021, doi:10.1088/1741-2560/10/3/036021 (2013).

68 Hiremath, S. V. et al. Human perception of electrical stimulation on the surface of somatosensory cortex. PLoS One 12, e0176020, doi:10.1371/journal.pone.0176020 (2017).

69 Murasugi, C. M., Salzman, C. D. & Newsome, W. T. Microstimulation in visual area MT: effects of varying pulse amplitude and frequency. J. Neurosci. 13, 1719–1729, doi:10.1523/jneurosci.13-04-01719.1993 (1993).

70 Jonas, J. et al. Intracerebral electrical stimulation of a face-selective area in the right inferior occipital cortex impairs individual face discrimination. Neuroimage 99, 487–497, doi:10.1016/j.neuroimage.2014.06.017 (2014).

71 Adab, H. Z. & Vogels, R. Perturbation of posterior inferior temporal cortical activity impairs coarse orientation discrimination. Cereb. Cortex 26, 3814–3827, doi:10.1093/cercor/bhv178 (2016).

72 Ringo, J. L. The medial temporal lobe in encoding, retention, retrieval and interhemispheric transfer of visual memory in primates. Exp. Brain Res. 96, 387–403, doi:10.1007/bf00234108 (1993).

73 Ringo, J. L. Brevity of processing in a mnemonic task. J. Neurophysiol. 73, 1712–1715, doi:10.1152/jn.1995.73.4.1712 (1995).

74 Lee, S.-H. et al. Activation of specific interneurons improves V1 feature selectivity and visual perception. Nature 488, 379–383, doi:10.1038/nature11312 (2012).

75 Song, Y.-H. et al. Somatostatin enhances visual processing and perception by suppressing excitatory inputs to parvalbumin-positive interneurons in V1. Sci. Adv. 6, eaaz0517, doi:10.1126/sciadv.aaz0517 (2020).

76 Atallah, B. V., Bruns, W., Carandini, M. & Scanziani, M. Parvalbumin-Expressing Interneurons Linearly Transform Cortical Responses to Visual Stimuli. Neuron 73, 159–170, doi:10.1016/j.neuron.2011.12.013 (2012).

77 Glickfeld, L. L., Histed, M. H. & Maunsell, J. H. R. Mouse primary visual cortex is used to detect both orientation and contrast changes. J. Neurosci. 33, 19416–19422, doi:10.1523/JNEUROSCI.3560-13.2013 (2013).

78 Lee, S.-H., Kwan, A. C. & Dan, Y. Interneuron subtypes and orientation tuning. Nature 508, E1–2, doi:10.1038/nature13128 (2014).

79 Reuter, J. H. Tilt discrimination in the mouse. Behav. Brain Res. 24, 81–84, doi:10.1016/0166-4328(87)90038-6 (1987).

80 Andermann, M. L., Kerlin, A. M. & Reid, C. Chronic cellular imaging of mouse visual cortex during operant behavior and passive viewing. Front. Cell. Neurosci. 4, doi:10.3389/fncel.2010.00003 (2010).

81 Ibrahim, L. A. et al. Bottom-up inputs are required for establishment of top-down connectivity onto cortical layer 1 neurogliaform cells. Neuron 109, 3473–3485.e3475, 10.1016/j.neuron.2021.08.004 (2021).

82 Tasic, B. et al. Adult mouse cortical cell taxonomy revealed by single cell transcriptomics. Nat. Neurosci. 19, 335–346, doi:10.1038/nn.4216 (2016).

83 Hartung, J., Schroeder, A., Péréz Vázquez, R. A., Poorthuis, R. B. & Letzkus, J. J. Layer 1 NDNF interneurons are specialized top-down master regulators of cortical circuits. Cell Rep. 43, 114212, doi:10.1016/j.celrep.2024.114212 (2024).

84 Tamás, G., Lorincz, A., Simon, A. & Szabadics, J. Identified sources and targets of slow inhibition in the neocortex. Science 299, 1902–1905, doi:10.1126/science.1082053 (2003).

85 Overstreet-Wadiche, L. & McBain, C. J. Neurogliaform cells in cortical circuits. Nat. Rev. Neurosci. 16, 458–468, doi:10.1038/nrn3969 (2015).

86 Douglas, R. J. & Martin, K. A. C. Neuronal circuits of the neocortex. Annu. Rev. Neurosci. 27, 419–451, doi:10.1146/annurev.neuro.27.070203.144152 (2004).

87 Miller, R. J. Presynaptic receptors. Annu. Rev. Pharmacol. Toxicol. 38, 201–227, doi:10.1146/annurev.pharmtox.38.1.201 (1998).

88 MacDermott, A. B., Role, L. W. & Siegelbaum, S. A. Presynaptic ionotropic receptors and the control of transmitter release. Annu. Rev. Neurosci. 22, 443–485, doi:10.1146/annurev.neuro.22.1.443 (1999).

89 Wang, L., Kloc, M., Maher, E., Erisir, A. & Maffei, A. Presynaptic GABAA receptors modulate thalamocortical inputs in layer 4 of rat V1. Cereb. Cortex 29, 921–936, doi:10.1093/cercor/bhx364 (2019).

90 Naumann, L. B. & Sprekeler, H. Presynaptic inhibition rapidly stabilises recurrent excitation in the face of plasticity. PLoS Comput. Biol. 16, e1008118, doi:10.1371/journal.pcbi.1008118 (2020).

91 Haley, M. S., Fontanini, A. & Maffei, A. Inhibitory gating of thalamocortical inputs onto rat gustatory insular cortex. J. Neurosci. 43, 7294–7306, doi:10.1523/JNEUROSCI.2255-22.2023 (2023).

92 Naumann, L. B., Hertäg, L., Müller, J., Letzkus, J. J. & Sprekeler, H. Layer-specific control of inhibition by NDNF interneurons. bioRxiv, doi:10.1101/2024.04.29.591728 (2024).

93 Poorthuis, R. B. et al. Rapid neuromodulation of layer 1 interneurons in human neocortex. Cell Rep. 23, 951–958, doi:10.1016/j.celrep.2018.03.111 (2018).

94 Huang, H. et al. Single pulse electrical stimulation in white matter modulates iEEG visual responses in human early visual cortex. bioRxiv, 2025.2005.2005.652264, doi:10.1101/2025.05.05.652264 (2025).

95 Beauchamp, M. S. et al. Dynamic Stimulation of Visual Cortex Produces Form Vision in Sighted and Blind Humans. Cell 181, 774–783.e775, doi:10.1016/j.cell.2020.04.033 (2020).

96 Halgren, A. S., Siegel, Z., Golden, R. & Bazhenov, M. Multielectrode cortical stimulation selectively induces unidirectional wave propagation of excitatory neuronal activity in biophysical neural model. J. Neurosci. 43, 2482–2496, doi:10.1523/JNEUROSCI.1784-21.2023 (2023).

97 Kumaravelu, K., Sombeck, J., Miller, L. E., Bensmaia, S. J. & Grill, W. M. Stoney vs. Histed: Quantifying the spatial effects of intracortical microstimulation. Brain Stimul. 15, 141–151, doi:10.1016/j.brs.2021.11.015 (2022).

98 Kumaravelu, K. et al. A comprehensive model-based framework for optimal design of biomimetic patterns of electrical stimulation for prosthetic sensation. J. Neural Eng. 17, 046045, doi:10.1088/1741-2552/abacd8 (2020).

99 Niell, C. M. & Stryker, M. P. Modulation of Visual Responses by Behavioral State in Mouse Visual Cortex. Neuron 65, 472–479, doi:10.1016/j.neuron.2010.01.033 (2010).

100 Zhong, J. et al. FACED 2.0 enables large-scale voltage and calcium imaging in vivo. bioRxiv, 2025.2003.2006.641784, doi:10.1101/2025.03.06.641784 (2025).

101 Ganji, M. et al. Selective formation of porous pt nanorods for highly electrochemically efficient neural electrode interfaces. Nano Lett. 19, 6244–6254, doi:10.1021/acs.nanolett.9b02296 (2019).

102 Tchoe, Y. et al. Human brain mapping with multithousand-channel PtNRGrids resolves spatiotemporal dynamics. Sci. Transl. Med. 14, eabj1441, doi:10.1126/scitranslmed.abj1441 (2022).

103 Fan, J. L. et al. High-speed volumetric two-photon fluorescence imaging of neurovascular dynamics. Nat. Commun. 11, doi:10.1038/s41467-020-19851-1 (2020).

104 Thunemann, M. et al. Deep 2-photon imaging and artifact-free optogenetics through transparent graphene microelectrode arrays. Nat. Commun. 9, 2035, doi:10.1038/s41467-018-04457-5 (2018).

105 Gage, G. J. et al. Surgical implantation of chronic neural electrodes for recording single unit activity and electrocorticographic signals. J. Vis. Exp., 3565, doi:10.3791/3565 (2012).

106 Chung, J. E. et al. Chronic implantation of multiple flexible polymer electrode arrays. J. Vis. Exp., 10.3791/59957, doi:10.3791/59957 (2019).

107 Lu, R. et al. Rapid mesoscale volumetric imaging of neural activity with synaptic resolution. Nat. Methods, doi:10.1038/s41592-020-0760-9 (2020).

108 Carandini, M. & Ferster, D. Membrane potential and firing rate in cat primary visual cortex. J. Neurosci. 20, 470–484 (2000).

